# Multiple coexisting pathways to synchronization shape seizure dynamics in a mesoscale mouse brain model

**DOI:** 10.64898/2026.06.08.730931

**Authors:** Nimish Kumar, Saurabh R. Gandhi

## Abstract

The computational study of epileptic seizure dynamics has primarily focused on the identification of seizure onset zones and propagation pathways. Here, we present a network dynamical model implemented on the empirically measured mesoscale mouse brain network that reveals previously unresolved organizational principles underlying seizure propagation. Rather than the conventional assumption of a single dominant pathway to synchronization, the model reveals multiple competing pathways to synchronization with distinct dynamical properties including transition propensity, recruitment speed, spatial coverage, synchronization stability and transition kinetics. Consequently, node ablation does not uniformly suppress synchronization across pathways, but instead selectively alters pathway occupancy, producing non-trivial alterations to seizure dynamics with potential implications for resection and network-targeted intervention strategies. Biologically, the olfactory and limbic sub-networks emerge as key mesoscale regulators of synchronization dynamics, with olfactory recruitment preferentially constraining global synchronization while limbic-driven pathways preferentially support seizure generalization. More broadly, these findings extend transient explosive synchronization theory by demonstrating that synchronization in biologically constrained networks may emerge through competing mesoscale recruitment programs rather than a single transition process. Together, these findings introduce a new conceptual framework for seizure propagation, suggesting that pathological synchronization emerges not through a single dominant route, but through competing mesoscale dynamical pathways whose accessibility depends on both network architecture and ongoing network state.

**Significance statement:** Epileptic seizure propagation is conventionally understood as progressing through a dominant pathway that recruits increasingly larger portions of the brain into pathological synchronization. Using a network dynamical model implemented on the empirical mesoscale mouse connectome, we show that seizure-like synchronization instead emerges through multiple competing pathways with distinct spatial and temporal characteristics. These pathways differ in their propensity for generalization, synchronization stability and sensitivity to node perturbation, such that network interventions selectively reshape pathway accessibility rather than uniformly suppressing seizure dynamics. Our findings introduce a new framework for understanding seizure propagation, identify mesoscale mechanisms linking network architecture to synchronization dynamics and suggest that competing synchronization pathways may represent an important organizing principle in complex brain networks.

**Graphical abstract:** 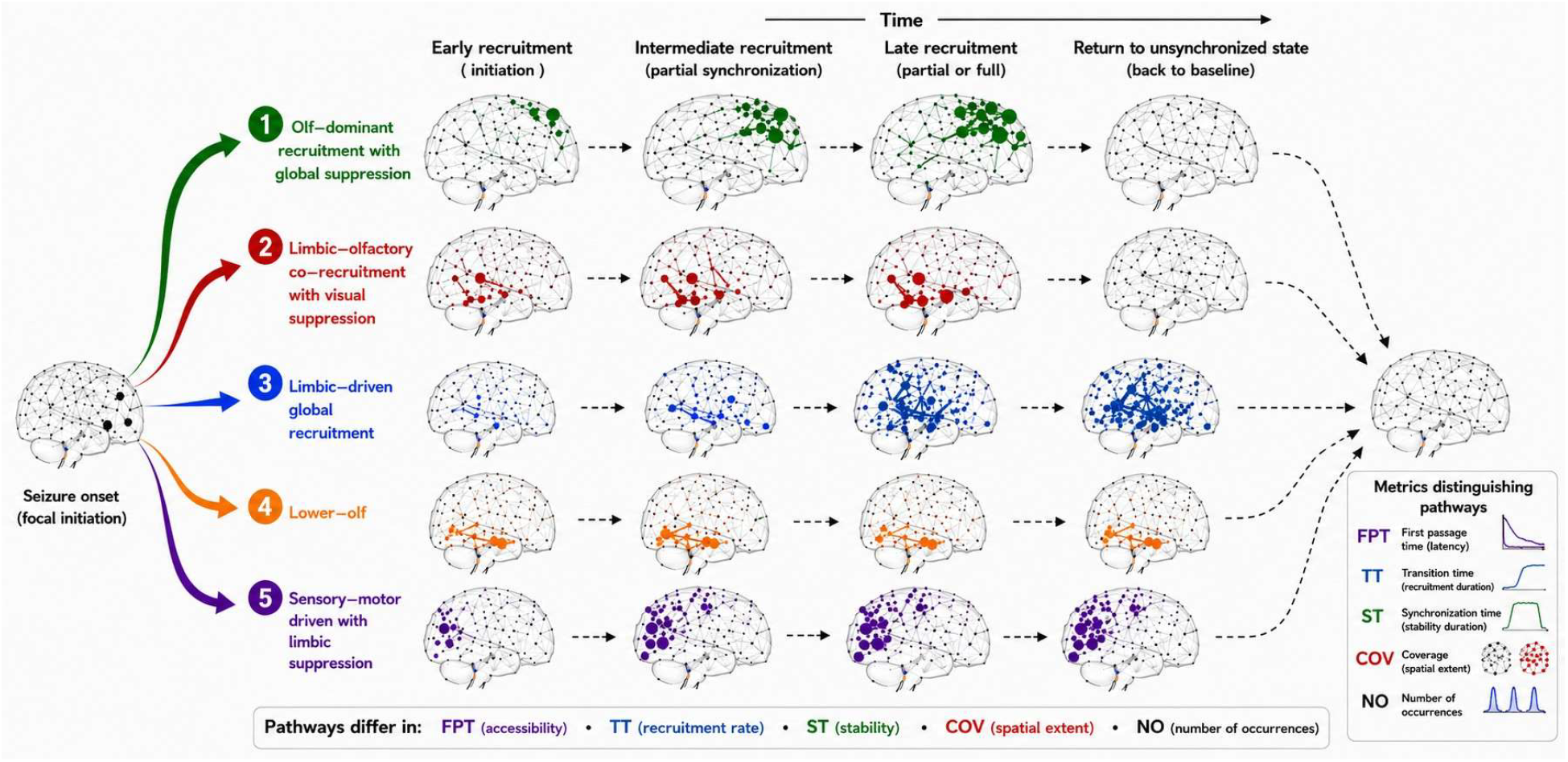

## Introduction

Seizures emerge through the recruitment of distributed brain regions into pathological synchronization, transforming initially localized activity into large-scale synchronized network dynamics (Olmi et al., 2019). Although substantial progress has been made in identifying seizure onset zones and propagation networks (Courson, 2025; Karoly et al., 2018), the dynamical principles governing how synchronization spreads through brain networks remain incompletely understood. Seizure propagation is often implicitly conceptualized as proceeding through a dominant synchronization pathway. However, the mammalian brain is a highly redundant and modular system capable of supporting multiple competing dynamical trajectories, raising the possibility that seizure generalization may emerge through multiple coexisting pathways to synchronization (Schroeder et al., 2022). Empirical investigation of such mesoscale dynamics remains challenging because of limitations in spatial sampling (Baud et al., 2021), motion artifacts during seizures (Vergult et al., 2007) and recording constraints (Miller et al., 2024; Parvizi and Kastner, 2018). Whole-brain network modeling informed by increasingly accurate connectomic measurements therefore provides a powerful framework for uncovering mesoscale mechanisms governing seizure propagation (Karoly et al., 2018).

Seizure propagation has been studied using network dynamical models spanning a wide range of model complexity and biological detail. While general principles of seizure dynamics have been elucidated using mean-field or biologically informed synthetic network models (Ersöz et al., 2020; Naze et al., 2015; Stefanescu et al., 2012), several recent studies have leveraged empirically measured connectivity in humans and animals to gain deeper mechanistic insights or make personalized predictions. For example, complex, biologically realistic models have been used to simulate seizure dynamics and generate patient-specific predictions for understanding epileptic networks and treatment strategies (Sun et al., 2024). Another extensively used family of phenomenological models is the Epileptor model, using which researchers have studied seizure initiation, propagation pathways and transitions between ictal and interictal states (Jirsa et al., 2014). On the other hand, minimalistic models such as those based on Kuramoto oscillator networks have also been used to explain empirical observations while identifying generic dynamical mechanisms underlying seizure propagation (Frolov and Hramov, 2021; Ranjan and Gandhi, 2024; Schmidt et al., 2014). By abstracting away biological complexity while retaining a few variables of interest, such models enable mechanistic insight with substantially reduced computational complexity.

While localization of seizure onset zones and propagation trajectory remains medically important (Rosenow and Lüders, 2001), seizure dynamics encompass additional features beyond where seizures begin and how they spread. Synchronization events may differ in their frequency, temporal stability, recruitment speed, spatial extent and composition of recruited communities, potentially reflecting distinct underlying pathways to synchronization. Distinguishing such pathways may have important implications for understanding seizure generalization and therapeutic intervention, as different synchronization routes may differ in their propensity for large-scale recruitment and their susceptibility to perturbation. Characterizing these properties may therefore provide a richer understanding of seizure propagation dynamics beyond conventional onset-zone or propagation-network descriptions. Here we quantify multiple such dynamical properties, including transition frequency, transition latency, synchronization stability and coverage (Fig. 1), and examine how they reveal competing mesoscale routes to synchronization.

**Fig. 1.**
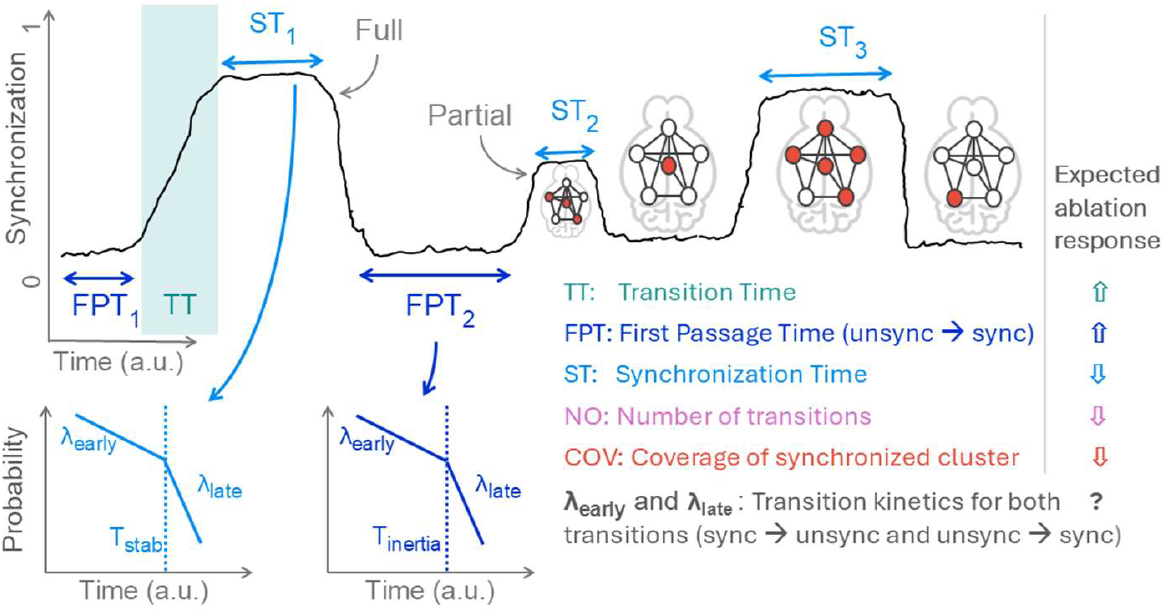
Dynamical characteristics of synchronization transitions quantified in this study and their hypothesized response to ablations. Coverage (COV): Percentage of network nodes participating in synchronization (green nodes). Ablation may reduce COV. **Transition Time (TT):** Duration required for the network to stabilize from transition onset to the synchronized state. Ablation may increase TT. **Number of Occurrences (NO):** Total number of synchronization transitions observed across simulations. Ablations may decrease NO. **Synchronization Time (ST):** Duration for which the network remains in the synchronized state following transition onset. Ablation may decrease ST. **First Passage Time (FPT):** Duration spent in the unsynchronized state before initiation of a synchronization transition. Ablation may increase FPT.

The theory of explosive synchronization, describing abrupt transitions from unsynchronized to synchronized states in complex networks, provides a broad framework for understanding seizure dynamics (Gómez-Gardeñes et al., 2011). Our previous work demonstrated how a minimal transient explosive synchronization (tES) model can recapitulate empirical seizure features while generating experimentally testable predictions (Ranjan and Gandhi, 2024). Explosive synchronization has been extensively studied in terms of synchronization onset and transition dynamics (Lee et al., 2025). Here, we further characterize competing synchronization pathways, their dynamical properties and their sensitivity to network node ablation by leveraging our tES framework implemented on the empirical mouse connectome. We further examine how structural and functional notions of node importance influence pathway recruitment and seizure propagation dynamics, thereby testing the hypothesis that seizure propagation emerges through multiple coexisting pathways to synchronization.

## Methods

### Model

Our model consists of *N* coupled sinusoidal oscillators representing nodes of the empirical brain network. Oscillator interactions follow a modified Kuramoto framework in which the coupling strength between oscillators is determined by the empirical structural connectivity matrix and is further modulated by two dynamic factors: (i) local synchronization levels within neighboring network regions and (ii) availability of a finite excitability resource (Eqn. 1 – 3) (Ranjan and Gandhi, 2024). Together, these mechanisms enable the network to exhibit transient explosive synchronization, characterized by abrupt transitions between unsynchronized and synchronized states followed by spontaneous desynchronization. The resulting dynamics provide a minimal mesoscale framework for studying seizure-like synchronization and propagation in large-scale brain networks.

### Brief description of the model, with details in (Ranjan and Gandhi, 2024)

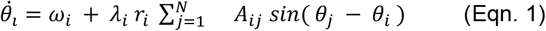

Here, *i* ∈ [1, *N*], *θ*_*i*_ and 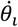 are instantaneous phase and angular velocity of the *i*^*th*^ oscillator, and *ω*_*i*_ is its natural frequency, uniformly distributed in [−1, 1]. *A*_*ij*_ encodes the adjacency matrix.

Interaction strengths are modulated by the local synchrony parameter,

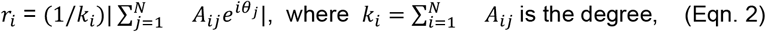

The interactions are also modulated by the availability of resources to individual nodes, *λ*_*i*_, whose dynamics is modeled as:

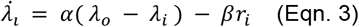

where the first term represents the recovery of excitability resources at a rate α, and the second term represents the local synchrony-dependent resource consumption at a rate β*r*_*i*_. β is the maximal consumption rate (when *r*_*i*_ = 1). The capacity of the resource reserve for each node is denoted by *λ*_*o*_(size of resource bath).

### Simulation details

Each network configuration is simulated 200 independent times for 40,000 epochs per simulation. Model parameters remain fixed across all simulations at α = 0.01, β = 0.002 and λ_0_ = 2.86. These parameters were chosen such that transitions from the unsynchronized to synchronized state are induced reliably in all networks. The global synchronization order parameter (Ranjan and Gandhi, 2024) is recorded throughout each simulation, while local synchronization measures are extracted within a temporal window spanning −3000 to +2000 time units relative to transition onset. Simulations were performed using Python version 3.10.19.

To characterize mesoscale recruitment dynamics during synchronization events, node community membership is tracked for every detected transition using the cluster lineage tracking framework (see methods, Ranjan and Gandhi, 2024). Community-level node participation is extracted within the transition-centered temporal window described above.

Simulations are performed on the mesoscale mouse brain connectome derived from the Allen Brain Atlas (Oh et al., 2014). The connectome is constructed using viral tracer experiments that quantify axonal projections between brain regions in healthy mice. The resulting network comprises 426 brain regions spanning both hemispheres and approximately 11,000 directed structural connections. Edge weights are normalized between 0 and 1 following previous work (Ranjan and Gandhi, 2024). This empirically derived mesoscale network serves as the structural substrate governing oscillator interactions in the transient explosive synchronization framework. Simulations performed on this intact connectome are referred to as the original Mesoscale Mouse Brain Network (**MMBN**) condition.

To evaluate the contribution of specific mesoscale regions, bilateral node ablations are performed by removing the corresponding nodes from both hemispheres of the connectome. Networks generated following node removal are referred to as ablated networks, by the name of the ablated node. All ablated networks are simulated using identical model parameters and simulation conditions. A total of 13 network configurations (one intact and twelve ablated) are analyzed.

### Choice of nodes for ablation

To compare structural and functional notions of node importance, we identify candidate nodes using both network topology and synchronization dynamics. Functional importance is quantified based on the median time spent by a node within the principal synchronization cluster across simulations. Nodes that remain synchronized for longer durations and possess a higher degree are hypothesized to exert greater influence on downstream recruitment and propagation dynamics.

Structural importance is quantified using network connectivity measures. Nodes with higher degree are expected to exert stronger influence on neighboring regions owing to their greater structural integration within the connectome. To identify nodes combining both structural and functional importance, we select the top six nodes exhibiting prolonged synchronization cluster participation together with high structural degree: ORB, PIR, BLA, RE, ENTI and PERI (Ranjan and Gandhi, 2024).

As controls, we additionally select nodes exhibiting dissociation between structural and functional properties, including nodes with high degree but low synchronization cluster participation (MRN), and nodes with low degree but high synchronization cluster participation (MOB, LSc). Finally, to assess structurally important regions independent of functional relevance, we calculate the rich-club coefficient (Heuvel and Sporns, 2011) across different degree thresholds. We normalize these values by comparing our graph to 20 randomized networks with the same degree distribution. A rich-club organization is confirmed wherever this normalized score is greater than 1, and the final hub elite is identified using the highest threshold that meets this condition, yielding LHA, MOs, PERI and MOp.

### Detection and classification of full and partial transitions

The global synchronization order parameter (Ranjan and Gandhi, 2024) is used to identify transitions between unsynchronized and synchronized states. To robustly detect synchronization transitions, we employ Uniform Manifold Approximation and Projection (UMAP) to obtain a low-dimensional representation of global order dynamics in running windows of length 400 time units. UMAP is first fitted on a representative simulation exhibiting three frequently observed conditions - unsynchronized, intermediate level of synchronization and high synchronization (the `skelarn 1.7.0` implementation of UMAP is used with `n_neighbours = 15`, `n_components = 2`, `min_dist = 0.1`, `metric = euclidean`) (Fig. 2A, B). The fitted embedding is subsequently used to project rolling windows of 400 consecutive global-order datapoints from all simulations into a two-dimensional embedding space. This representation separates synchronized (including partially synchronized) and unsynchronized states based on a threshold on the first dimension (Fig. 2C-F), allowing individual timepoints to be classified accordingly.

**Fig. 2.**
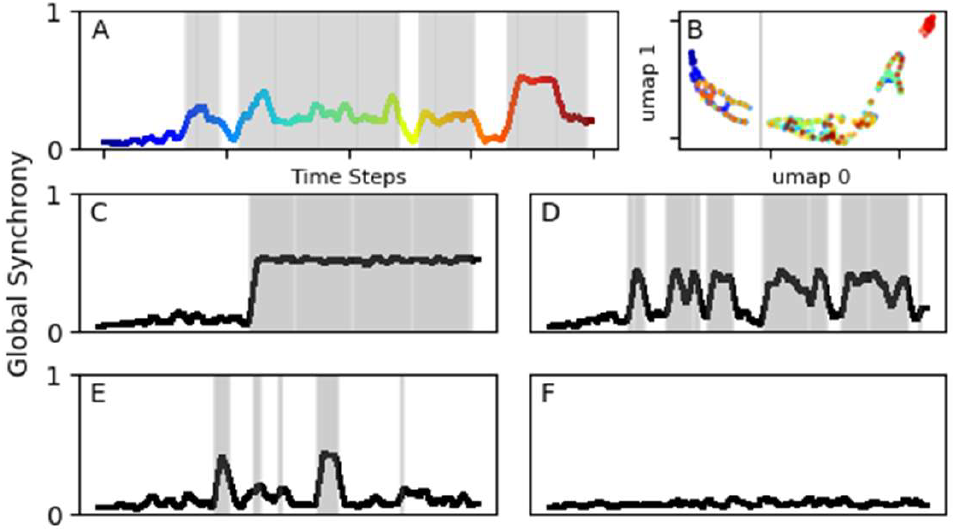
Automated detection and identification of synchronization transitions using UMAP. **(A)** The synchronization time course from MMBN, exhibiting both partial and full synchronization events, which was used to train the UMAP embedding model. **(B)** Two-dimensional UMAP projection of rolling-window global-order dynamics from the time course shown in **A**, separating transition and non-transition states. **(C–F)** Example synchronization time courses illustrating automated transition identification across simulations. Grey shaded regions indicate detected synchronization transitions. Example simulations shown in **C–F** contain 4, 6, 5 and 0 detected transitions, respectively, including full and partial ones.

Transition onset is identified using two criteria. First, the global order parameter must exceed 0.2. Second, transition-associated embeddings must persist for at least four consecutive timepoints. The transition endpoint is defined as the first subsequent timepoint at which the global order parameter falls below 0.3.

Detected synchronization events are further classified as partial or full transitions using the node coverage metric. Coverage distributions within each network configuration exhibit bimodal structure, corresponding to low-coverage and high-coverage synchronization states. The minimum of the Kernel Density Estimate separating these two modes is used as a network-specific threshold to classify individual synchronization events as partial or full transitions.

### Cluster tracking

The Synchronization Cluster Tracking Algorithm (SCTA) employs a two-step procedure to quantify node participation during synchronization dynamics (Ranjan and Gandhi, 2024). First, synchronization clusters are generated by initializing from a seed node and iteratively expanding cluster membership until local synchrony falls below a fixed threshold. Second, synchronization clusters are tracked progressively through time to construct cluster lineages, enabling quantification of temporal recruitment dynamics and node participation during synchronization events.

### Node communities

Node communities are identified on the empirical connectome using the Louvain community detection algorithm, following the framework introduced previously (Ranjan and Gandhi, 2024). Community detection partitions the network into eight mesoscale communities corresponding broadly to orbital (ORB), sensory–motor (MO), visual/auditory (VIS), olfactory (OLF), hypothalamic (HTH), hippocampal (HIPP), midbrain (MID) and hindbrain (HIND) subnetworks. These mesoscale communities serve as the basis for community-level recruitment analysis during synchronization transitions. Community participation profiles are subsequently used to characterize how different functional systems contribute to seizure propagation dynamics and distinct synchronization pathways supported by the network.

### Definitions of metrics (Fig. 1)

#### Transition Time (TT)

Transition Time quantifies the duration required for the network to stabilize into a synchronized state following transition initiation. Transition onset is identified using the UMAP-based procedure described above. From the detected transition onset, the slope of the global order parameter is computed using a rolling window of 200 time units. The endpoint of transition dynamics is defined as the point at which the slope approaches zero, indicating stabilization of synchronization. Transition Time is then computed as the interval between transition onset and this stabilization point.

#### First Passage Time (FPT)

First Passage Time quantifies the latency required for the network to transition from the unsynchronized state to synchronization onset. FPT is computed as the elapsed simulation time from simulation initiation or the end of previous transition to the transition onset identified using the UMAP-based transition detection framework.

#### Number of Occurrences (NO)

Number of Occurrences quantifies transition frequency and is defined as the total number of synchronization transitions observed across 200 simulations. Each transition onset identified using the UMAP framework contributes one occurrence.

#### Synchronization Time (ST)

Synchronization Time quantifies the duration for which the network remains in the synchronized state during a transition event. ST is computed as the interval between transition onset and transition endpoint identified using the UMAP-based procedure.

#### Coverage (COV)

Coverage quantifies the spatial extent of synchronization and is defined as the percentage of nodes participating in synchronization during a transition event. Coverage is computed by averaging node-level synchronization participation between transition onset and transition endpoint.

### Statistical analysis

For each of the metric TT, FPT, ST and COV, we compare ablated networks with MMBN using Mann-Whitney U-test with Holm-Bonferroni correction method. The Poisson mean test with Holm-Bonferroni correction method is used to compare NO of MMBN to ablated networks (Fig. 3A). Two-tailed Wald Z-test for the difference between two independent slope estimates with Holm-Bonferroni correction is used to compare significance of transition rates, *T*_*inertia*_ and stabilisation time (Fig. 5I, J) with respect to MMBN. Pearson correlation is used to find the correlation of parameters with betweenness centrality (Fig. 6). To determine statistically significant differences in transition dynamics across the identified subtypes, we performed pairwise comparisons using the non-parametric Mann-Whitney U-test. To control the family-wise error rate associated with multiple comparisons, significance thresholds were sequentially adjusted using the Holm-Bonferroni method, where a corrected p < 0.05 was considered statistically significant (Fig. 5B, 5F). To determine differences in State Transition (ST) rates between network configurations and across time points (Fig. 5H), two-sided non-parametric Mann-Whitney U tests were carried out, again with multiple comparison correction.

**Fig. 3.**
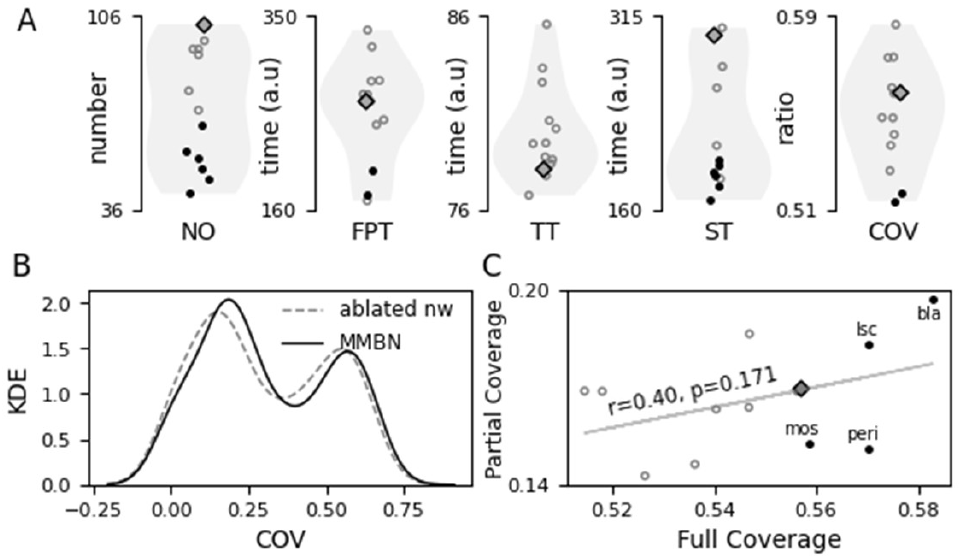
Effects of node ablation on synchronization dynamics across network configurations. **(A)** Effect of node removal on Number of Occurrences (NO), First Passage Time (FPT), Transition Time (TT), Synchronization Time (ST) and Coverage (COV). Y-axis units correspond to time units for FPT, TT and ST, while NO denotes the number of transitions observed across 200 simulations. The intact MMBN condition is shown as a diamond, while filled circles indicate networks exhibiting significant differences relative to MMBN. Mean values across all observed transitions for 13 networks are shown. **(B)** Distribution of transition coverage values (Kernel Density Estimate) for the intact MMBN and an example ablated network (ORB), illustrating the bimodal structure used to classify synchronization events. **(C)** Relationship between coverage for partial and full synchronization transitions for the different network.

## Results

### Differential impact of ablation across dynamical parameters

We begin our analysis by evaluating the impact of node removal on the parameters identified earlier (Fig. 3A). Broadly, the number of transitions to the synchronized state (NO), time spent in the synchronized state (ST) as well as the mean coverage of the synchronized state (COV) decrease in ablated networks compared to the original mesoscale mouse brain network (MMBN). The transition time (TT), i.e. the time for the transition to complete once initiated, increases for most ablated networks. These observations are consistent with the intuition that removing nodes might make transitions to the synchronized state difficult (NO decreases), slow (TT increases), unstable (ST decreases) or incomplete (COV decreases). The first passage time (FPT) to the synchronized state does not follow a predictable pattern, increasing or decreasing for the different ablated networks.

Interestingly, in all our simulations, irrespective of the identity of the ablated node, we observe a bimodal distribution of coverage across transitions – in one subset of transitions, the synchronized cluster of nodes covers a substantial fraction of the total number of nodes (henceforth called a full transition) while for remaining transitions, the synchronized cluster covers a smaller fraction of nodes (henceforth called partial transitions) (Fig. 3B). Node removal significantly decreases the coverage of both full and partial transitions in general, although we observe some exceptions to this (Fig. 3C). Moreover, although partial transitions generally outnumber full transitions, the number of full and partial transitions is not correlated across networks (Fig. 3C). Together these two observations suggest differences across networks in terms of both the probability of transition to synchronized state, as well as disassociation between full and partial synchronization.

### Synchronized states exhibit different subtypes based on their node composition

Next we investigate the node-level composition of the synchronized states for individual transitions to synchrony. We decompose the empirical mouse brain network into eight structure-based node communities using the Louvain algorithm, and compute the average participation of each community in the synchronized cluster of nodes, giving us an 8-feature vector for each transition. We then apply UMAP-based projection from this 8D to a 2D space followed by clustering to identify five transition subtypes based on node participation in the synchronized state, observable across all networks (Fig. 4A, Fig. S4.1,S4.2).

**Fig. 4.**
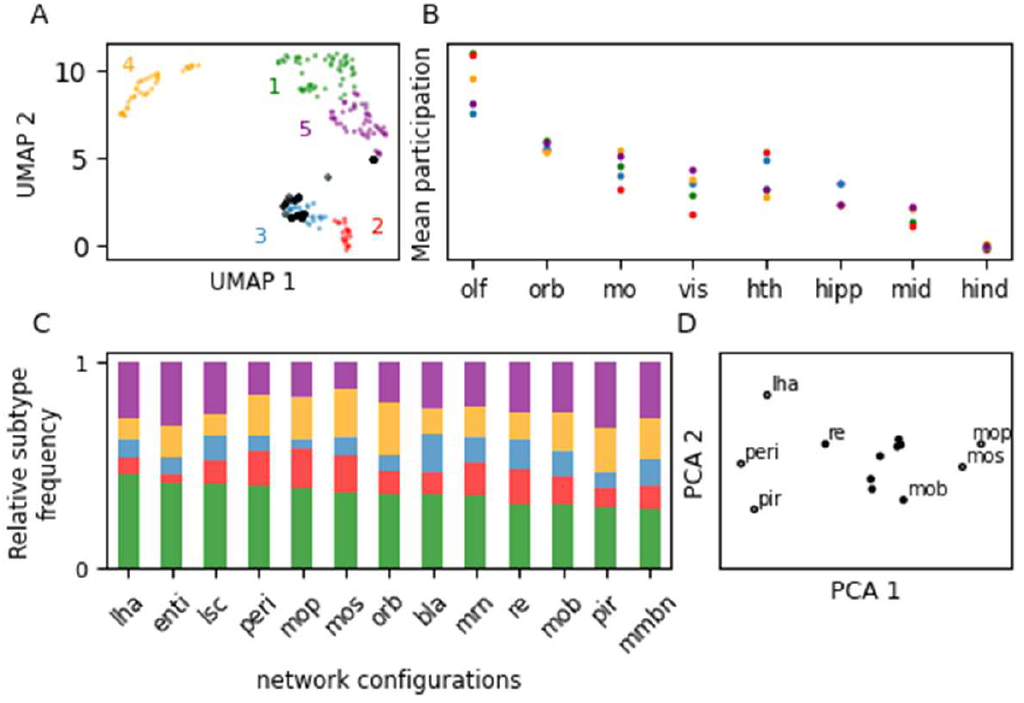
Community-level composition of synchronization subtypes and their modulation by node ablation. **(A)** Community participation profiles for synchronization transitions in the intact Mesoscale Mouse Brain Network (MMBN), projected using UMAP into two dimensions from eight and clustered, revealing five distinct synchronization subtypes. Black circles denote full synchronization transitions, which predominantly localize to subtype 3, and occasionally to subtype 5. **(B)** Community participation profiles across synchronization subtypes, highlighting strong olfactory (OLF) recruitment across all subtypes. **(C)** Relative occupancy of synchronization subtypes across network configurations (tick label denotes ablated node). Colored bars indicate the proportion of transitions assigned to each subtype within a given network. **(D)** Community participation profiles across all 13 network configurations projected into two dimensions using PCA. Networks involving LHA, PERI and PIR ablations (as also MOB, MOS, MOP) exhibit similar community recruitment profiles, suggesting related effects on synchronization pathway composition.

To validate our non-linear dimensionality reduction and clustering, and understand the transition subtype properties, we use the subtype identities from above to perform supervised classification on the original features using the random forest algorithm. The resulting SHAP analysis (Lundberg et al., 2020) elucidates the driving factors for each of the five identified subtypes as follows (Fig. 4B, Fig. S4.3): (1) **olfactory-dominant recruitment** with global suppression; (2) **limbic–olfactory co-recruitment** with visual suppression; (3) **limbic-driven global recruitment**; (4) **lower-olfactory participation**; (5) **sensory–motor driven** with limbic suppression. The subtypes are thus primarily driven by the relative participation levels of three community families in the synchronized clusters: limbic, sensory-motor and olfactory.

Full transitions are predominantly observed in only two of the five transition subtypes identified above (subtypes 3 – limbic driven global and 5 – sensory-motor with relative limbic suppression, Fig. 4A). Subtype 3 especially shows few partial transitions, which are otherwise spread across all five subtypes, although appearing dominantly in subtype 1 (OLF recruitment). To summarize, observe several subtypes of partial transitions differing in community participation, and a separate class of full transitions in all simulated networks.

The relative subtype frequencies differ across the networks we simulated (Fig. 4C). Subtype 1 is the most prevalent and stable in all networks. The frequencies of subtypes 2 and 4 are correlated, and anticorrelated with subtype 5 (Fig. S4.4). Networks with ablated PERI, MOs, MOp, RE and MRN exhibit a preference for these subtypes (2 and 4; away from 5 i.e. high limbic-olfactory and low sensory-motor participation) while the opposite (high sensory-motor participation) holds for the remaining networks. This is consistent with the location of the ablated nodes - for example, it is not surprising that the ablation of the motor area (MOp) reduces the occurrence of sensory-motor synchronization (subtype 5) (Fig. 4C). The varying relative frequency of subtypes observed in different networks leads us to hypothesize the existence of multiple coexisting competing pathways to synchronization, corresponding to the different subtypes, with certain pathways getting subdued upon ablation of specific nodes.

To understand how the community participation profile differs across different ablated networks, we associated each network with a 40-feature vector comprising the participation level of each community (8 in number) in each transition subtype (5 in number), and projected it in 2D with PCA (Fig. 4D). We find networks with ablations of LHA, PERI or PIR form a distinct cluster, as do MOs and MOp compared to the remaining networks. The distinctions are primarily driven by the level of OLF community participation in subtype 3 (anticorrelated with sensory-motor participation): the ablation of motor areas reduced the recruitment of motor regions in subtype 3 transitions and increased OLF recruitment; and vice versa for areas LHA, PERI and PIR (Fig. S4.5, S4.6).

### DIfferent transition subtypes reflect different synchronization pathways

A major impact of node removal one might expect is the blocking of certain synchronization pathways. In our analysis, this would be reflected in a change in the relative number of transitions across different subtypes, or a change in the conversion ratio to full transition.

Considering the subtypes to reflect distinct synchronization pathways, we hypothesize that the relative numbers of full to partial transitions in a subtype might be a good way to quantify the efficacy of the pathway to go all the way up to full synchronization. For example, in the original MMBN, we observe that subtype 3 (limbic-driven) is a highly successful pathway to full brain synchronization, while subtype 5 (sensory-motor driven) shows some but not very high conversion (Fig. 5A). On the other hand, subtypes 1, 2 and 4, all having increased OLF participation, never culminate in the fully synchronized state, hinting at an important role played by the OLF community, which upon recruitment seems to somehow impede further full synchronization.

**Fig. 5:**
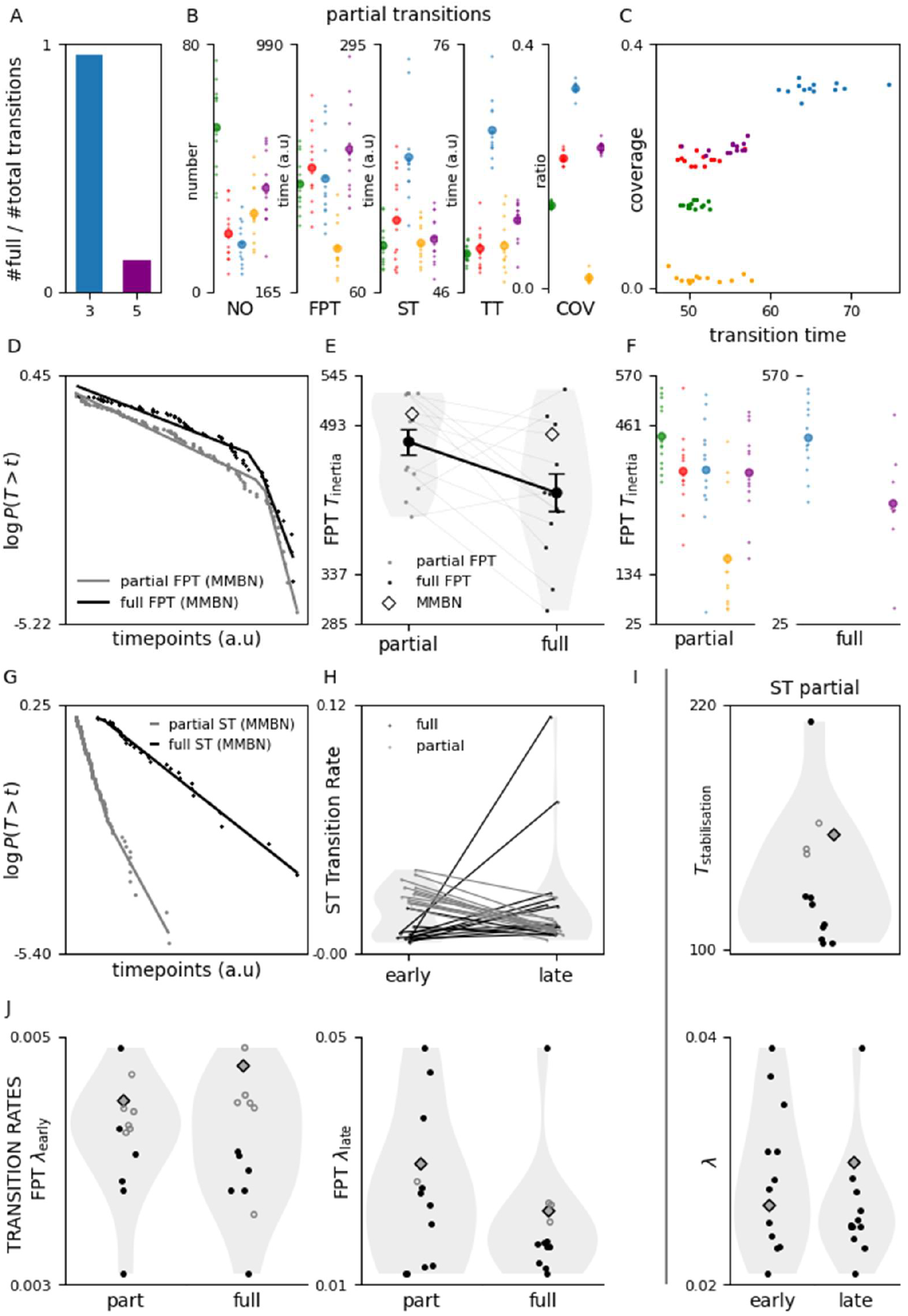
Subtype characteristics reflect distinct synchronization pathways. **(A)** Likelihood of conversion from partial to full transition. **(B)** Average characteristic values for different subtypes (partial transitions only). Individual networks shown as small dots and mean across networks shown as large dots. **(C)** Relationship between coverage and transition time for partial transitions in the five subtypes. **(D)** Piecewise exponential distribution of first passage time for both full (black) and partial (gray) transitions shown for MMBN (see SI for other networks). **(E)** FPT inertia for partial and full transitions (individual networks in light gray and mean in black). **(F)** FPT inertia for individual subtypes, partial (left) and full (right) transitions. **(G)** Piecewise exponential distribution of time in synchronized state (ST) for full and partial transitions (MMBN). **(H)** ST transition rates increase for partial (gray) and decrease for full (black) transitions. **(I)** Stabilization time for partial transitions (top) and corresponding early and late transition rates (bottom). **(J)** Transition rates for FPT (early and late) for full and partial transitions.

We further characterize these different transition subtypes, based on the dynamical parameters identified earlier (Fig. 5B, see SI Table 5.1 for significance of differences). The maximum number of partial transitions end up in subtype 1, followed by subtype 5. Subtypes 2, 3 and 4 show the least number of partial transitions, of which 2 is low because most of them convert to full transitions. In apparent contradiction, although subtype 4 has relatively fewer partial transitions, it shows a significantly lower first passage time to synchrony. It also shows the least coverage. These observations may be reconciled as follows: in an initial time window immediately after returning to the unsynchronized state, the networks are susceptible to enter into a short (least COV) pathway represented by subtype 4 (reduced FPT). This window is short, leading to a reduced overall NO irrespective of the low FPT. Other pathways, activated at later times, are more likely to result in a partially synchronized state.

Subtypes 2 and 3 tend to be more stable in the partially synchronized state, reflected in a higher ST. However, while subtype 2 is able to convert this stable partial sync to a full sync, subtype 3 never successfully proceeds to full synchronization. Coverage of partial transitions exhibits the highest variation across subtypes – highest for subtype 3, and lowest for subtype 4. Transition time is highest for subtype 3, followed by subtypes 5, 4, 2 and 1, consistent with the coverage – larger coverages require longer time to achieve. However, the transition times are not proportional to the coverage, with subtypes 1 and 2 showing much slower transitions compared to their coverage. These subtype dependent differences are consistent with the subtypes representing distinct independent pathways (Fig. 5C).

### Frequency and stability of the synchronized state across full and partial transitions

To better understand the dynamics of conversion from partial to full transitions, we investigate the transition probabilities into and out of the different synchronized states. We define the first passage time to synchrony as the time from onset of the unsynchronized state (either simulation onset or just after returning to the unsynchronized state) to onset of the full or partial synchronized state. We find that the FPT for both the full and partial transitions follow a piecewise exponential distribution. At short times post onset of the unsynchronized state, the likelihood of transitioning back to synchrony decreases with a slow exponent (Fig. 5D, S5.1,5.2), reflecting a Poisson process for transitioning into the synchronized state at some rate. At longer times however, this likelihood decreases at a faster exponential rate, corresponding to a higher Poisson transition rate. This reflects inertia in the network to retain the unsynchronized state for some duration before it switches to a state with a higher propensity to transition to synchrony. This inertia likely reflects the time required for the network to recover energy resources that are rapidly depleted in the synchronized state.

If the pathways to partial and full synchronization were shared, i.e. all full transitions followed the pattern from unsynchronized → partial sync → full sync, we would expect the switching time to be lesser or at best the same for partial compared to full transitions. Yet, we observe the switching time to be relatively shorter for transitions to full synchrony (∼400 time units post previous desynchronization) compared to transitions to partial synchrony (∼500 time units post previous desynchronization) (Fig. 5E; paired t-test t = 2.43, p = 0.032). This is possible only if the two pathways are independent, with the partial synchronization pathways having more inertia compared to the full synchronization pathways. This is further corroborated by the switching time for individual transition subtypes: subtype 5 transitions have a lower inertia for full transitions compared to partial transitions. Subtypes 1, 2, 3 and 5 partial transitions as well as subtype 3 full transition show no significant difference in inertia (Fig. 5F, S5.6, S5.7).

Transitions from the synchronized to unsynchronized state also follow a piecewise exponential ST (synchronization time, the analog of FPT for the sync → unsync transition) (Fig. 5g, S5.3). Full transitions remain stable for long durations, often up to the end of the simulation run. Therefore, we are able to obtain very limited data for full transition ST, resulting in poor quality of piecewise exponential fits (Fig. S5.4). However, the general trend we observe is that the desynchronization is slower at first, and increases later, due to the rapid resource depletion in the synchronised state (Fig. 5H; Mann-Whitney U test; u = 131, p = 0.0015).

In contrast, the transition rates back to the unsynchronized state from partial synchronization are larger at shorter times, and decrease at longer times (Fig. 5H, Mann-Whitney U test; u = 5, p < 0.001). This suggests a non-trivial tension between resource depletion (which is slower in the partially synchronised state) pushing for desynchronization and further recruitment pushing for expansion. Thus, the network seems to stabilize in the partially synchronized state after a short duration, with a high fallback rate before stabilization and a reduced rate after.

The transition rate to the synchronized state (full or partial) is lower in ablated networks (Fig. 5J, 5 / 12 networks for partial and 6 / 12 for full transitions are significant for early times, 8 / 12 partial and 7 / 12 full at later times), reflecting the increased difficulty in synchronization (both at early times post-desynchronization as well as later times). Surprisingly though, ablation generally leads to a decrease in the FPT inertia for both full and partial transitions (T test; t = 2.43, p = 0.03 ; Fig. 5E), i.e. the switch to higher transition rates happens sooner in ablated networks. This might be consistent with decreased coverage in ablated networks, which in turn result in lower energy depletion and thus lower recovery times (inertia).

Ablation does not alter the stability of the partially synchronized state in terms of the initial rate of desynchronization (Fig. 5I). Surprisingly, though, they stabilize sooner in the partially synchronized state (8 / 12 networks show significance, SI Table 5.1), and after stabilization, have a lower rate of desynchronization compared to the original MMBN (11 / 12 networks show significance, SI Table 5.1). Partial transitions are thus surprisingly more stable in ablated networks compared to the MMBN. Full transitions, although having sparse data, show the opposite trend (Fig. S5.4): they take longer to stabilize in ablated networks, and desynchronize faster even after stabilization compared to the MMBN. Therefore the overall synchronized time is higher on average for MMBN compared to ablated networks, consistent with our observation in Fig. 3B.

These observations reflect the extremely non-trivial dynamics of transition to synchronization in the mouse brain network.

### The role of structurally and functionally important nodes

Lastly, we compare the impact of the removal of structurally as opposed to functionally important nodes in the mouse brain network (Fig. 6). Only the number of full and partial transitions is significantly altered upon ablation: 4 / 6 functionally (and only 1 / 4 structurally) important nodes show significant decrease in partial transitions (Fig. 6A). 4 / 6 functional and 3 / 4 structurally important nodes show a significant decrease in full transitions. The number of full transitions, transition time and conversion ratio to full synchrony are all significantly negatively correlated with the degree centrality of the ablated node (Fig. 6A, rightmost column). We thus see a predominant effect of functionally important nodes on partial transitions, while both structurally and functionally important nodes have a significant effect on full transitions.

**Fig. 6.**
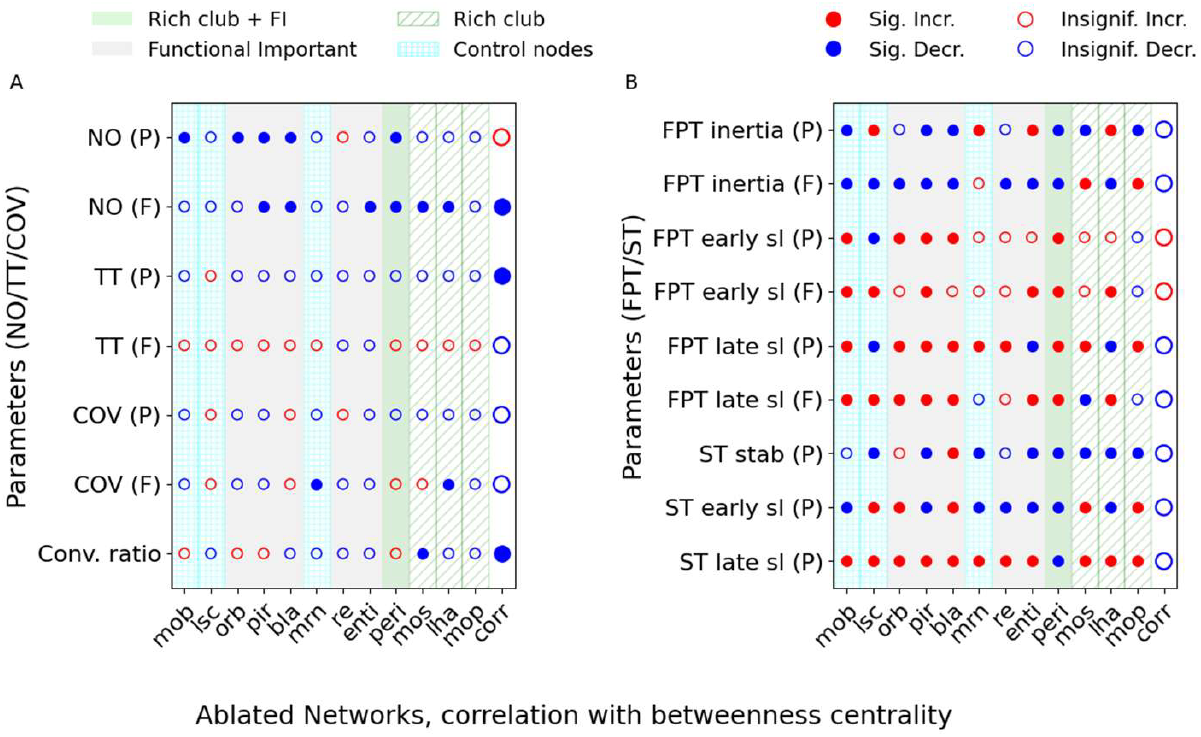
Effects of ablating structurally and functionally important nodes on synchronization dynamics. Each row represents a dynamical metric and each column represents an ablated node. Nodes are arranged along the x-axis in increasing order of betweenness centrality. Metric values indicate deviations relative to the intact Mesoscale Mouse Brain Network (MMBN) condition. The final column shows the correlation between betweenness centrality and the corresponding metric. Significant deviations from MMBN (p<0.05) are denoted by filled circles, while non-significant changes are shown as open circles. Red indicates positive deviations and blue indicates negative deviations relative to MMBN.

Nearly all kinetic parameters are significantly impacted by node ablation, irrespective of the nature of the node, although no significant correlation is observed with the node centrality (Fig. 6B).

## Discussion

In this study, we use a transient explosive synchronization (tES) framework implemented on an empirically derived mesoscale mouse brain connectome to investigate the network mechanisms underlying seizure-like synchronization dynamics. Rather than treating synchronization as a single homogeneous process, we characterize it as a repertoire of distinct pathways with different recruitment patterns, temporal characteristics and sensitivities to node ablation. Moving beyond conventional analyses focused primarily on seizure onset zones or single propagation trajectories (Emerson et al., 2020; Proix et al., 2014; Sinha et al., 2017), our approach combines cluster tracking, community-level participation analysis and systematic node ablation to identify synchronization subtypes reproducible across network configurations and examine how specific mesoscale regions bias the network toward particular synchronization pathways. Together, these analyses reveal that seizure-like synchronization emerges through multiple coexisting mesoscale pathways rather than a single dominant route.

The synchronization subtypes identified in our analysis broadly organize into two families of recruitment pathways. Subtypes 1, 2 and 4 exhibit stronger olfactory participation and predominantly manifest as partial transitions, suggesting locally stable yet globally constrained synchronization modes. Subtype 1, the most prevalent across networks, represents a readily accessible pathway that rarely progresses to global recruitment. Subtype 4 represents a low-energy pathway that is available in the energy depleted state immediately after a synchronization event, although it is weakly sustained. Subtype 2 combines coordinated limbic–olfactory recruitment with prolonged synchronization stability, yet still fails to convert into full synchronization.

In contrast, subtypes 3 and 5 represent large-scale recruitment pathways associated with higher coverage and substantially greater proportions of full transitions. Subtype 3, driven by the limbic network, emerges as the most efficient pathway to global synchronization, whereas subtype 5 preferentially recruits sensory–motor regions and supports full synchronization less consistently. Together, these findings suggest that the identified subtypes correspond to qualitatively distinct mesoscale recruitment pathways rather than simple variations in synchronization strength.

Importantly, node ablation does not uniformly suppress synchronization across all pathways. Instead, ablations selectively alter the relative occupancy of different pathways. For example, motor-area ablations reduce the prevalence of sensory–motor dominated transitions, whereas ablations involving PERI, PIR or LHA preferentially reshape olfactory participation profiles. Node removal therefore alters the landscape of accessible synchronization pathways rather than merely weakening overall synchronizability. These observations suggest that mesoscale regions influence seizure dynamics not simply by increasing or decreasing synchronization propensity, but by biasing the network toward specific classes of recruitment pathways.

The distinct coexisting pathway hypothesis is further supported by the transition kinetics of full and partial synchronization, which are inconsistent with a simple serial progression from unsynchronized to partially synchronized to fully synchronized states. Instead, the shorter switching times observed for full transitions suggest that full and partial synchronization arise through independent dynamical channels. In this framework, synchronization is not a single attractor approached through different trajectories, but rather a repertoire of metastable pathways whose accessibility depends on both network structure and ongoing dynamical state.

The repeated involvement of olfactory and limbic communities across synchronization subtypes has important biological implications. Regions such as the piriform cortex and associated paralimbic structures have long been implicated in seizure susceptibility and propagation, particularly in temporal and limbic epilepsies (Galovic et al., 2019; Vismer et al., 2015). In our model, increased olfactory participation is consistently associated with partial synchronization states and reduced conversion to full synchronization, suggesting that olfactory circuitry supports locally persistent yet globally constrained recruitment dynamics. This observation is consistent with the low seizure threshold of the rodent piriform cortex and its proposed role in gating seizure generalization (Tao et al., 2026). In contrast, limbic recruitment appears context dependent. Although limbic–olfactory co-recruitment remains associated with partial synchronization, limbic-dominant recruitment emerges as the most effective pathway to widespread synchronization, suggesting that limbic circuits participate in both constrained and globally propagating synchronization modes depending on the broader mesoscale recruitment pattern. The strong influence of PERI and PIR ablations on subtype composition further suggests that these regions shape the balance between local and global synchronization pathways. Although our model does not establish physiological mechanisms directly, the consistency of these observations with known epileptogenic properties of piriform and limbic circuitry (Vaughan and Jackson, 2014) suggests that the identified synchronization subtypes may reflect biologically meaningful modes of seizure propagation.

Our findings also highlight an important distinction between structural and functional notions of node importance. Structurally central nodes preferentially influence full transitions and large-scale synchronization dynamics, whereas functionally important nodes—defined by sustained participation in synchronization clusters – affect both full and partial transitions. This divergence suggests that graph-theoretic centrality alone may be insufficient to capture the dynamical role of nodes in seizure propagation (Laiou et al., 2019). Highly connected hubs may facilitate large-scale integration and stabilization of globally synchronized states, while not being very relevant in local synchronization motifs. Nodes that remain persistently engaged within synchronization clusters, however, seem to support both local and global synchronization. These observations reinforce the idea that seizure propagation depends not only on static anatomical connectivity, but also on dynamically evolving recruitment patterns distributed across the network (Olmi et al., 2019). Consequently, functional participation profiles may provide complementary information to conventional structural metrics when identifying critical regions involved in seizure progression.

Current surgical approaches in epilepsy focus primarily on identifying and removing the seizure onset zone (Beniczky et al., 2025; Mazhit et al., 2026). However, our results suggest that perturbing a structural or functional node can induce non-trivial network-level reconfigurations rather than simply suppressing synchronization globally. While ablation of structurally important hubs reduces large-scale synchronization and seizure generalization, it also promotes faster stabilization of localized partial synchronization pathways. This raises the possibility that interventions suppressing generalized seizure propagation may nevertheless preserve—or unmask—alternative localized synchronization routes within the remaining network. These observations support the need for a broader network-level framework for surgical planning that accounts not only for seizure onset regions, but also for latent competing pathways that may become dynamically accessible following resection or neuromodulation (Khambhati et al., 2016).

More broadly, these results extend the conceptual scope of transient explosive synchronization models in brain networks. Most prior work on explosive synchronization focuses on whether synchronization occurs, the conditions required for abrupt transitions, or the hysteretic properties of the synchronized state (Bayani et al., 2022). In contrast, our findings suggest that synchronization in biologically constrained networks may be fundamentally multi-route rather than binary. The coexistence of multiple transient synchronization pathways indicates that explosive synchronization can give rise to heterogeneous metastable states rather than a single globally synchronized regime. Partial synchronization, in particular, emerges not merely as an incomplete transition state, but as a dynamically stable mesoscale configuration with its own temporal structure and pathway dependence. Similar pathway-dependent collective dynamics have been proposed in systems ranging from resting-state brain activity to cardiac fibrillation (Carrick et al., 2021; Courtiol et al., 2020). Together, these observations suggest that pathway competition may represent an important but underexplored dimension of synchronization phenomena in complex networks.

In conclusion, this study introduces a new perspective on seizure propagation by demonstrating that synchronization in the mesoscale mammalian brain emerges through multiple coexisting and competing pathways rather than a single dominant propagation route. The model generates specific experimentally testable predictions regarding pathway-selective perturbations in brain networks. In particular, our findings predict that targeted perturbation of limbic, olfactory or sensory–motor regions should not uniformly suppress seizure-like synchronization, but instead selectively bias the brain away from specific synchronization pathways while preserving or favoring others. Testing these predictions using large-scale electrophysiological or optical recordings together with targeted perturbation experiments could establish whether the identified synchronization subtypes correspond to biologically separable modes of seizure propagation in vivo. Understanding these pathways, rather than only identifying seizure onset regions, may therefore offer a richer framework for interpreting and controlling pathological synchronization in the brain, while also opening new avenues for network-centric and pathway-specific therapeutic strategies.

## Acknowledgements

SRG was supported by the Science and Engineering Research Board (SERB), Award ID: SRG/2023/000595.

## Author contributions

NK and SRG conceived the study, designed the model and computational framework, analyzed the data, and wrote the manuscript.

## Data availability

The code and parameters that have provided the results presented here are available on GitHub at https://github.com/csndl-iitd/srg-tes-epilepsy-models.

## References

1. Baud, M.O., Schindler, K., Rao, V.R., 2021. Under-sampling in epilepsy: Limitations of conventional EEG. Clinical Neurophysiology Practice 6, 41–49. 10.1016/j.cnp.2020.12.002

2. Bayani, A., Jafari, S., Azarnoush, H., 2022. Explosive synchronization: From synthetic to real-world networks. Chinese Phys. B 31, 020504. 10.1088/1674-1056/ac3cb0

3. Beniczky, S., Trinka, E., Wirrell, E., Abdulla, F., Al Baradie, R., Alonso Vanegas, M., Auvin, S., Singh, M.B., Blumenfeld, H., Bogacz Fressola, A., Caraballo, R., Carreno, M., Cendes, F., Charway, A., Cook, M., Craiu, D., Ezeala-Adikaibe, B., Frauscher, B., French, J., Gule, M.V., Higurashi, N., Ikeda, A., Jansen, F.E., Jobst, B., Kahane, P., Kishk, N., Khoo, C.S., Vinayan, K.P., Lagae, L., Lim, K.-S., Lizcano, A., McGonigal, A., Perez-Gosiengfiao, K.T., Ryvlin, P., Specchio, N., Sperling, M.R., Stefan, H., Tatum, W., Tripathi, M., Yacubian, E.M., Wiebe, S., Wilmshurst, J., Zhou, D., Cross, J.H., 2025. Updated classification of epileptic seizures: Position paper of the International League Against Epilepsy. Epilepsia 66, 1804–1823. 10.1111/epi.18338

4. Carrick, R.T., Benson, B.E., Bates, O.R.J., Spector, P.S., 2021. Competitive Drivers of Atrial Fibrillation: The Interplay Between Focal Drivers and Multiwavelet Reentry. Front. Physiol. 12. 10.3389/fphys.2021.633643

5. Courson, J., 2025. Neural network structure, onset and propagation of epileptic seizures (Theses). CY Cergy Paris Université ; University of Warwick (Royaume-Uni).

6. Courtiol, J., Guye, M., Bartolomei, F., Petkoski, S., Jirsa, V.K., 2020. Dynamical Mechanisms of Interictal Resting-State Functional Connectivity in Epilepsy. J. Neurosci. 40, 5572–5588. 10.1523/JNEUROSCI.0905-19.2020

7. Emerson, J., Afelin, A., Sliupas, V., Fink, C.G., 2020. Identifying Influential Nodes in a Network Model of Epilepsy. J Nonlinear Sci 30, 2283–2308. 10.1007/s00332-019-09545-4

8. Ersöz, E.K., Modolo, J., Bartolomei, F., Wendling, F., 2020. Neural mass modeling of slow-fast dynamics of seizure initiation and abortion. PLOS Computational Biology 16, e1008430. 10.1371/journal.pcbi.1008430

9. Frolov, N., Hramov, A., 2021. Extreme synchronization events in a Kuramoto model: The interplay between resource constraints and explosive transitions. Chaos: An Interdisciplinary Journal of Nonlinear Science 31, 063103. 10.1063/5.0055156

10. Galovic, M., Baudracco, I., Wright-Goff, E., Pillajo, G., Nachev, P., Wandschneider, B., Woermann, F., Thompson, P., Baxendale, S., McEvoy, A.W., Nowell, M., Mancini, M., Vos, S.B., Winston, G.P., Sparks, R., Prados, F., Miserocchi, A., de Tisi, J., Van Graan, L.A., Rodionov, R., Wu, C., Alizadeh, M., Kozlowski, L., Sharan, A.D., Kini, L.G., Davis, K.A., Litt, B., Ourselin, S., Moshé, S.L., Sander, J.W.A., Löscher, W., Duncan, J.S., Koepp, M.J., 2019. Association of Piriform Cortex Resection With Surgical Outcomes in Patients With Temporal Lobe Epilepsy. JAMA Neurol 76, 690–700. 10.1001/jamaneurol.2019.0204

11. Gómez-Gardeñes, J., Gómez, S., Arenas, A., Moreno, Y., 2011. Explosive Synchronization Transitions in Scale-Free Networks. Phys. Rev. Lett. 106, 128701. 10.1103/PhysRevLett.106.128701

12. Heuvel, M.P. van den, Sporns, O., 2011. Rich-Club Organization of the Human Connectome. J. Neurosci. 31, 15775–15786. 10.1523/JNEUROSCI.3539-11.2011

13. Jirsa, V.K., Stacey, W.C., Quilichini, P.P., Ivanov, A.I., Bernard, C., 2014. On the nature of seizure dynamics. Brain 137, 2210–2230. 10.1093/brain/awu133

14. Karoly, P.J., Kuhlmann, L., Soudry, D., Grayden, D.B., Cook, M.J., Freestone, D.R., 2018. Seizure pathways: A model-based investigation. PLOS Computational Biology 14, e1006403. 10.1371/journal.pcbi.1006403

15. Khambhati, A.N., Davis, K.A., Lucas, T.H., Litt, B., Bassett, D.S., 2016. Virtual Cortical Resection Reveals Push-Pull Network Control Preceding Seizure Evolution. Neuron 91, 1170–1182. 10.1016/j.neuron.2016.07.039

16. Laiou, P., Avramidis, E., Lopes, M.A., Abela, E., Müller, M., Akman, O.E., Richardson, M.P., Rummel, C., Schindler, K., Goodfellow, M., 2019. Quantification and Selection of Ictogenic Zones in Epilepsy Surgery. Front. Neurol. 10. 10.3389/fneur.2019.01045

17. Lee, S., Kuklinski, L.J., Timme, M., 2025. Extreme synchronization transitions. Nat Commun 16, 4505. 10.1038/s41467-025-59729-8

18. Lundberg, S.M., Erion, G., Chen, H., DeGrave, A., Prutkin, J.M., Nair, B., Katz, R., Himmelfarb, J., Bansal, N., Lee, S.-I., 2020. From local explanations to global understanding with explainable AI for trees. Nat Mach Intell 2, 56–67. 10.1038/s42256-019-0138-9

19. Mazhit, A., Akbay, B., Trofimov, A., Karapina, O., Duysenbi, S., Tokay, T., 2026. Epileptogenesis and Epilepsy Treatment: Advances in Mechanistic Understanding, Therapeutic Approaches, and Future Perspectives. Int J Mol Sci 27, 1175. 10.3390/ijms27031175

20. Miller, K.R., Barnard, S., Juarez-Colunga, E., French, J.A., Pellinen, J., 2024. Long-term seizure diary tracking habits in clinical studies: Evidence from the Human Epilepsy Project. Epilepsy Research 203, 107379. 10.1016/j.eplepsyres.2024.107379

21. Naze, S., Bernard, C., Jirsa, V., 2015. Computational Modeling of Seizure Dynamics Using Coupled Neuronal Networks: Factors Shaping Epileptiform Activity. PLOS Computational Biology 11, e1004209. 10.1371/journal.pcbi.1004209

22. Oh, S.W., Harris, J.A., Ng, L., Winslow, B., Cain, N., Mihalas, S., Wang, Q., Lau, C., Kuan, L., Henry, A.M., Mortrud, M.T., Ouellette, B., Nguyen, T.N., Sorensen, S.A., Slaughterbeck, C.R., Wakeman, W., Li, Y., Feng, D., Ho, A., Nicholas, E., Hirokawa, K.E., Bohn, P., Joines, K.M., Peng, H., Hawrylycz, M.J., Phillips, J.W., Hohmann, J.G., Wohnoutka, P., Gerfen, C.R., Koch, C., Bernard, A., Dang, C., Jones, A.R., Zeng, H., 2014. A mesoscale connectome of the mouse brain. Nature 508, 207–214. 10.1038/nature13186

23. Olmi, S., Petkoski, S., Guye, M., Bartolomei, F., Jirsa, V., 2019. Controlling seizure propagation in large-scale brain networks. PLOS Computational Biology 15, e1006805. 10.1371/journal.pcbi.1006805

24. Parvizi, J., Kastner, S., 2018. Promises and limitations of human intracranial electroencephalography. Nat Neurosci 21, 474–483. 10.1038/s41593-018-0108-2

25. Proix, T., Bartolomei, F., Chauvel, P., Bernard, C., Jirsa, V.K., 2014. Permittivity Coupling across Brain Regions Determines Seizure Recruitment in Partial Epilepsy. J. Neurosci. 34, 15009–15021. 10.1523/JNEUROSCI.1570-14.2014

26. Ranjan, A., Gandhi, S.R., 2024. Propagation of transient explosive synchronization in a mesoscale mouse brain network model of epilepsy. Network Neuroscience 8, 883–901. 10.1162/netn_a_00379

27. Rosenow, F., Lüders, H., 2001. Presurgical evaluation of epilepsy. Brain 124, 1683–1700. 10.1093/brain/124.9.1683

28. Schmidt, H., Petkov, G., Richardson, M.P., Terry, J.R., 2014. Dynamics on Networks: The Role of Local Dynamics and Global Networks on the Emergence of Hypersynchronous Neural Activity. PLOS Computational Biology 10, e1003947. 10.1371/journal.pcbi.1003947

29. Schroeder, G.M., Chowdhury, F.A., Cook, M.J., Diehl, B., Duncan, J.S., Karoly, P.J., Taylor, P.N., Wang, Y., 2022. Multiple mechanisms shape the relationship between pathway and duration of focal seizures. Brain Commun 4, fcac173. 10.1093/braincomms/fcac173

30. Sinha, N., Dauwels, J., Kaiser, M., Cash, S.S., Brandon Westover, M., Wang, Y., Taylor, P.N., 2017. Predicting neurosurgical outcomes in focal epilepsy patients using computational modelling. Brain 140, 319–332. 10.1093/brain/aww299

31. Stefanescu, R.A., Shivakeshavan, R.G., Talathi, S.S., 2012. Computational models of epilepsy. Seizure 21, 748–759. 10.1016/j.seizure.2012.08.012

32. Sun, J., Niu, Y., Wang, C., Dong, Y., Wang, B., Wei, J., Xiang, J., Ma, J., 2024. Exploring the propagation pathway in individual patients with epilepsy: A stepwise effective connection approach. Biomedical Signal Processing and Control 90, 105811. 10.1016/j.bspc.2023.105811

33. Tao, Y., Zhao, Y., Zhong, W., Zhang, J., Zhu, H., Zhu, X., Wang, Z., Wang, N., Yang, L., Xu, F., Wu, R., 2026. Piriform seizures mediated by the piriform-entorhino-dentate circuit induce brain-wide functional reorganization in mice. PLOS Biology 24, e3003577. 10.1371/journal.pbio.3003577

34. Vaughan, D.N., Jackson, G.D., 2014. The Piriform Cortex and Human Focal Epilepsy. Front. Neurol. 5. 10.3389/fneur.2014.00259

35. Vergult, A., De Clercq, W., Palmini, A., Vanrumste, B., Dupont, P., Van Huffel, S., Van Paesschen, W., 2007. Improving the Interpretation of Ictal Scalp EEG: BSS–CCA Algorithm for Muscle Artifact Removal. Epilepsia 48, 950–958. 10.1111/j.1528-1167.2007.01031.x

36. Vismer, M.S., Forcelli, P.A., Skopin, M.D., Gale, K., Koubeissi, M.Z., 2015. The piriform, perirhinal, and entorhinal cortex in seizure generation. Front. Neural Circuits 9. 10.3389/fncir.2015.00027

