## Supplementary Information for "Multiple coexisting pathways to synchronization shape seizure dynamics in a mesoscale mouse brain model"

Fig. S4.1: Five Transition subtypes based on node participation

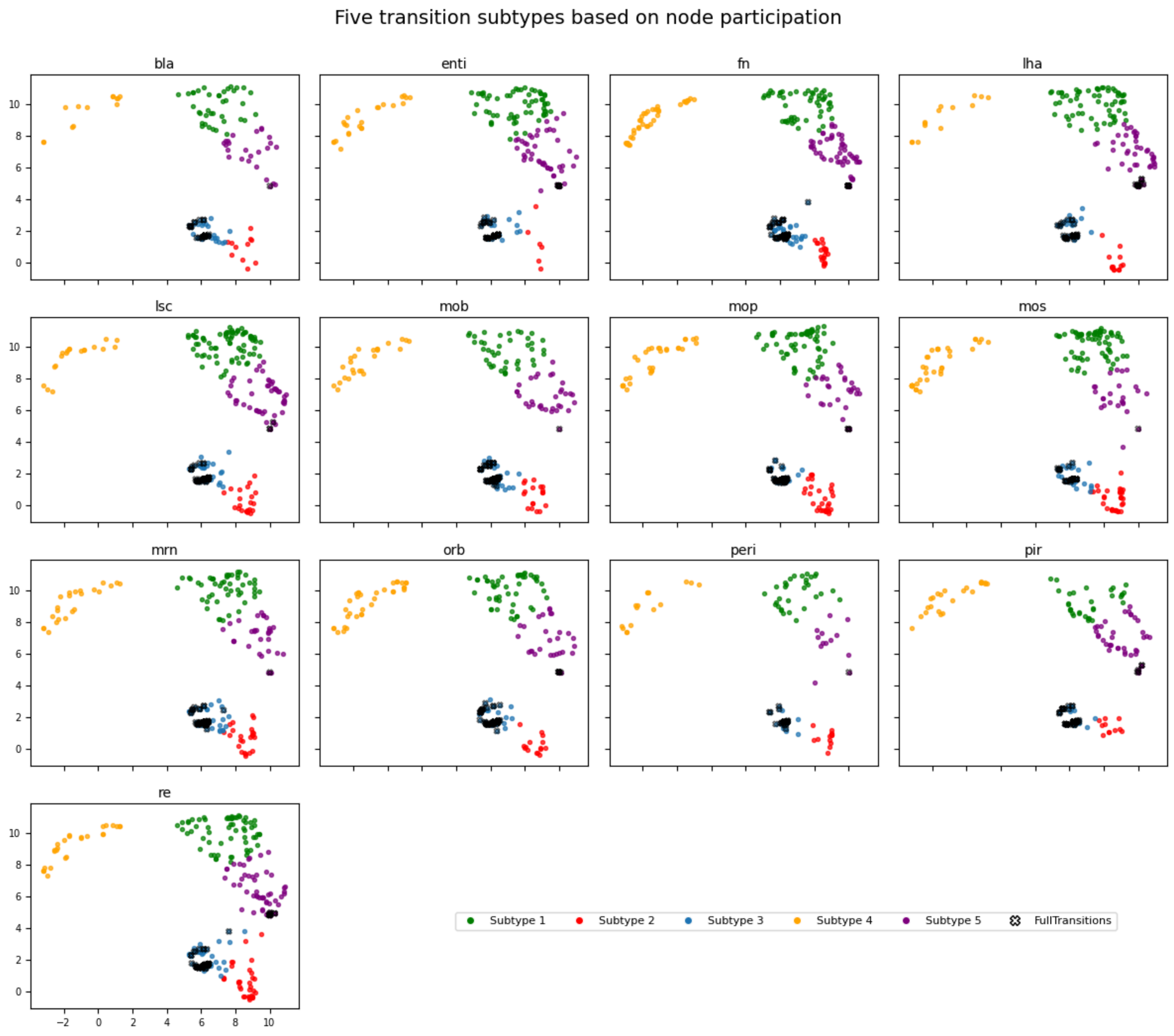


8 feature vector community participation is reduced to 2 dimensions using UMAP. Further clustered to give 5 subtypes indicating 5 kinds of seizure propagation pathways

Fig.S4.2: AIC / BIC

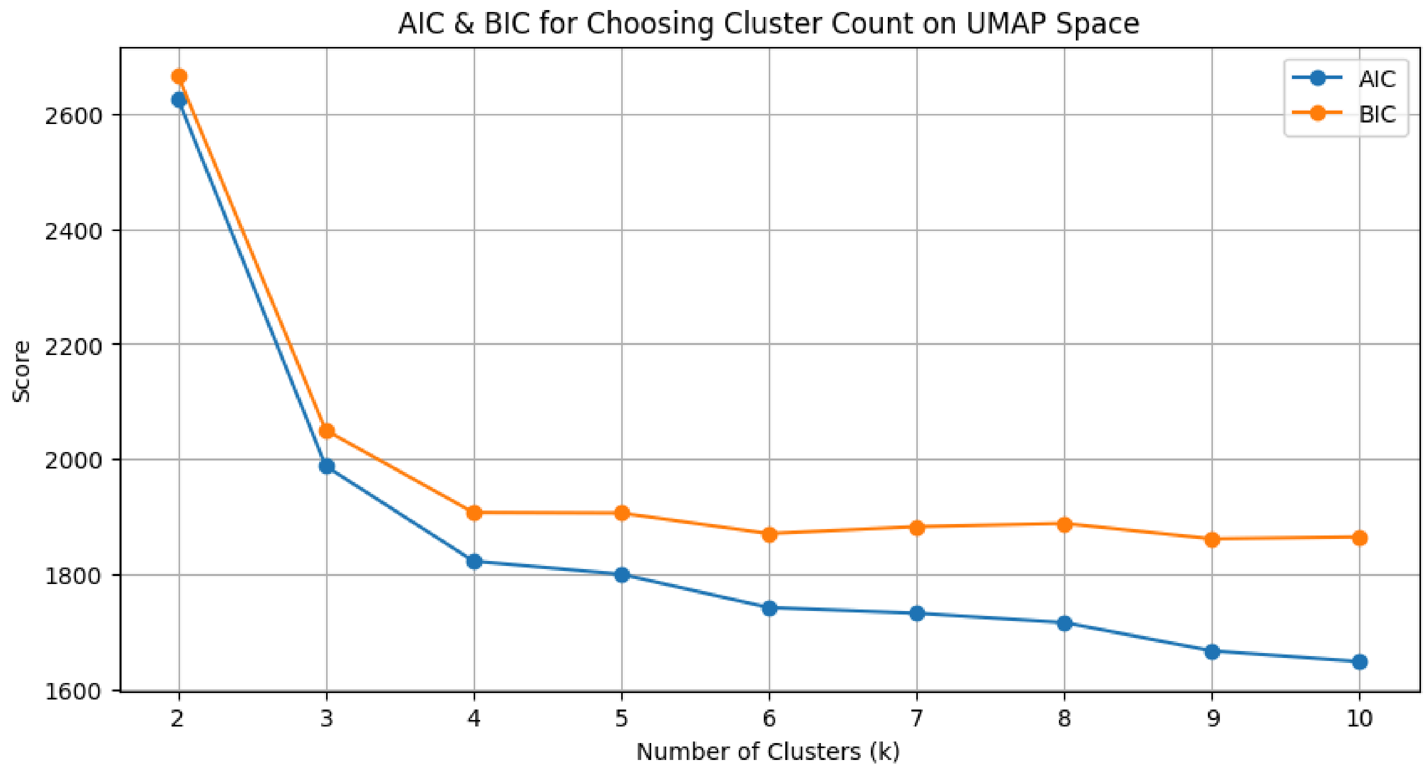


AIC/BIC performed on reduced MMBN community participation indicates 5 clusters to be optimum for clusterisation

Fig. S4.3: SHAP analysis

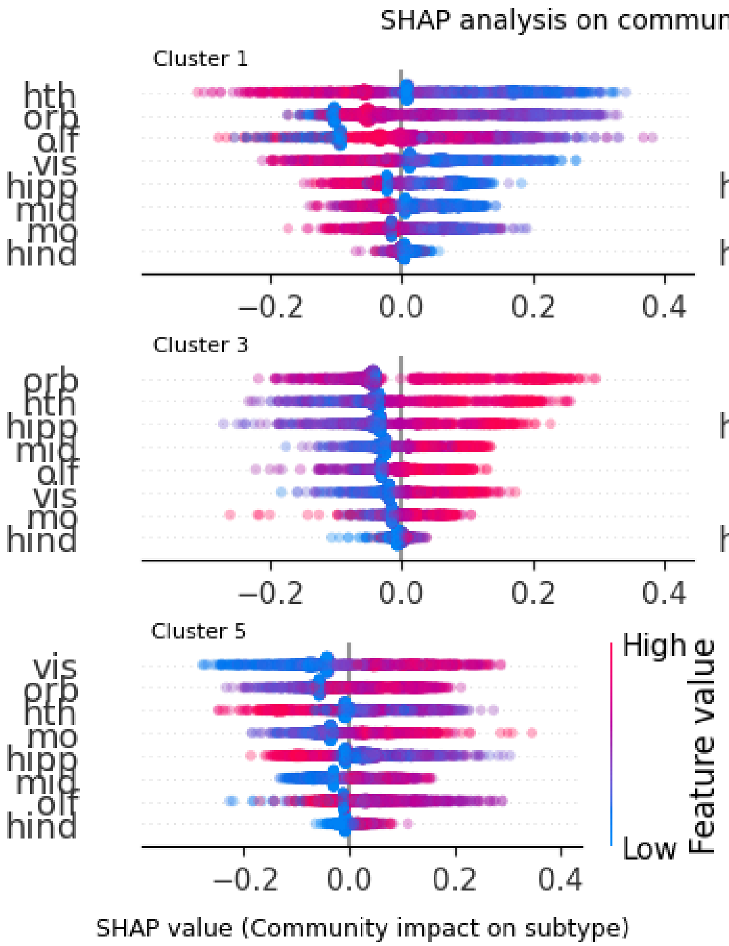


SHAP done on reduced data of all networks to show effect of community on specific clusters

Fig. S4.4: Subtypes preference to networks in general

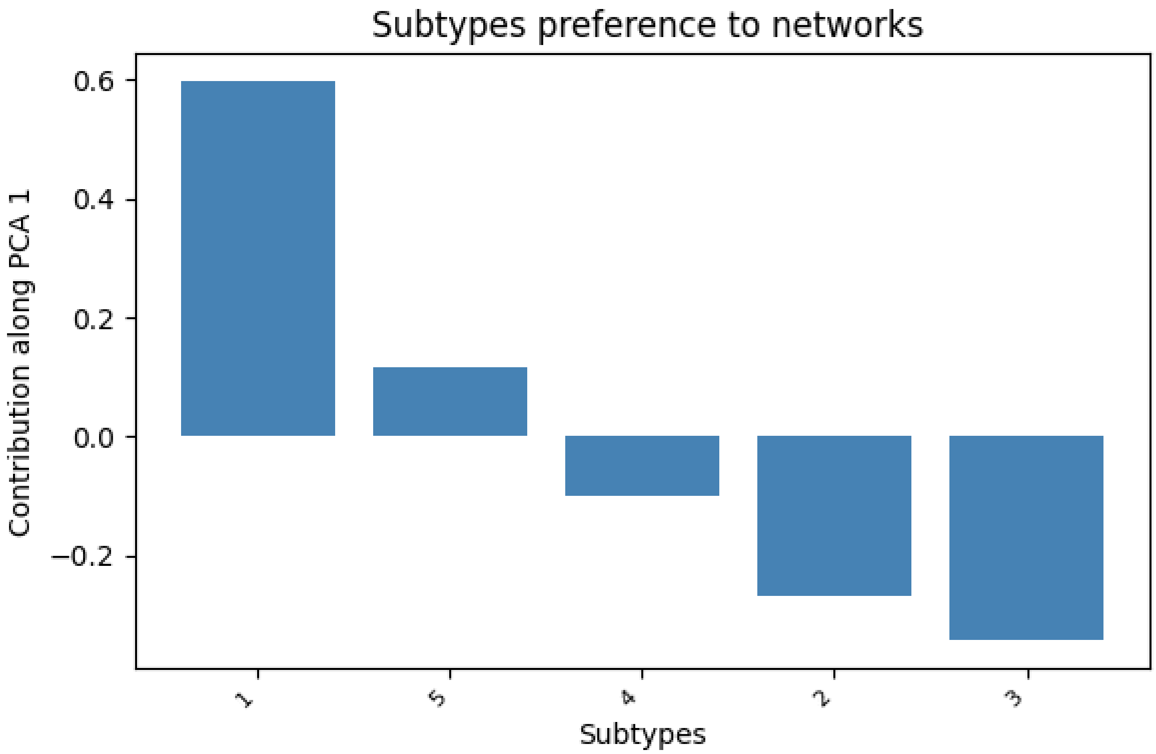


PCA done to show transition of which subtypes are more observed in general

Fig. S4.5: Subtype specific community preference across all the networks

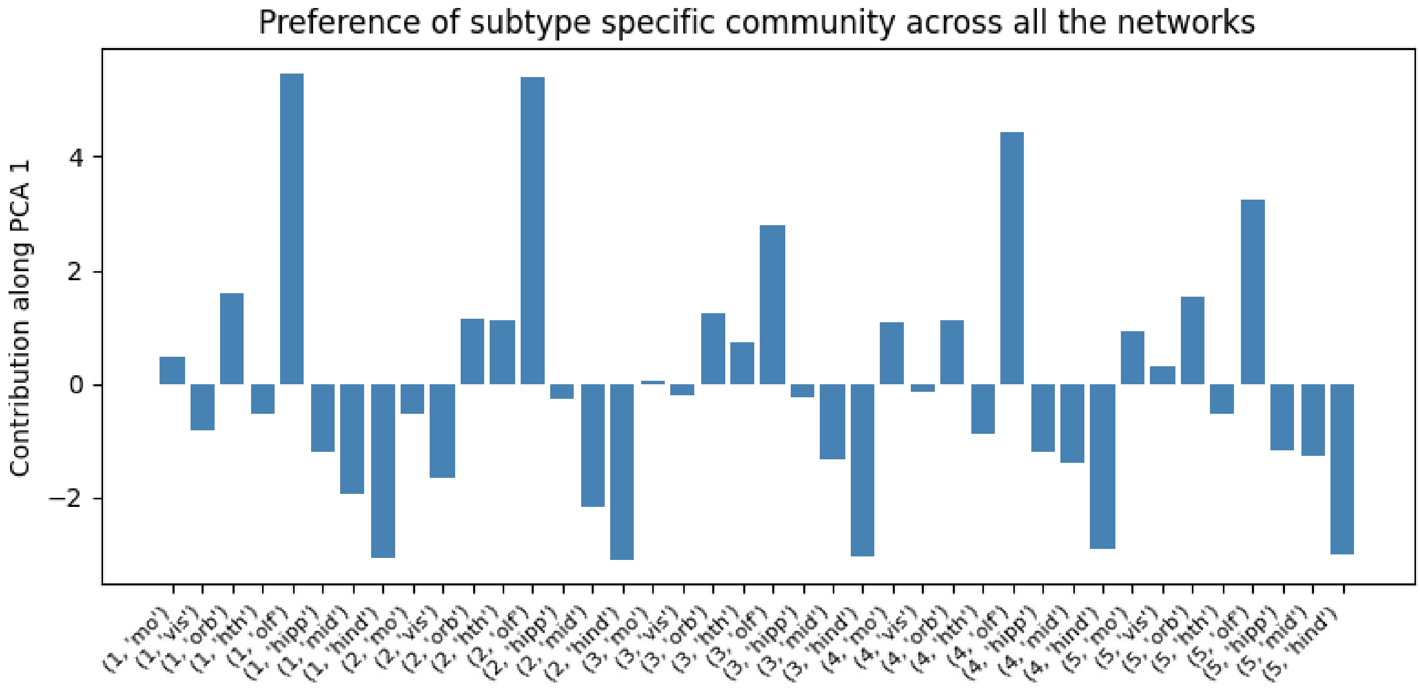


PCA done to show which subtype and community majorly participates across all transitions

Fig. S4.6

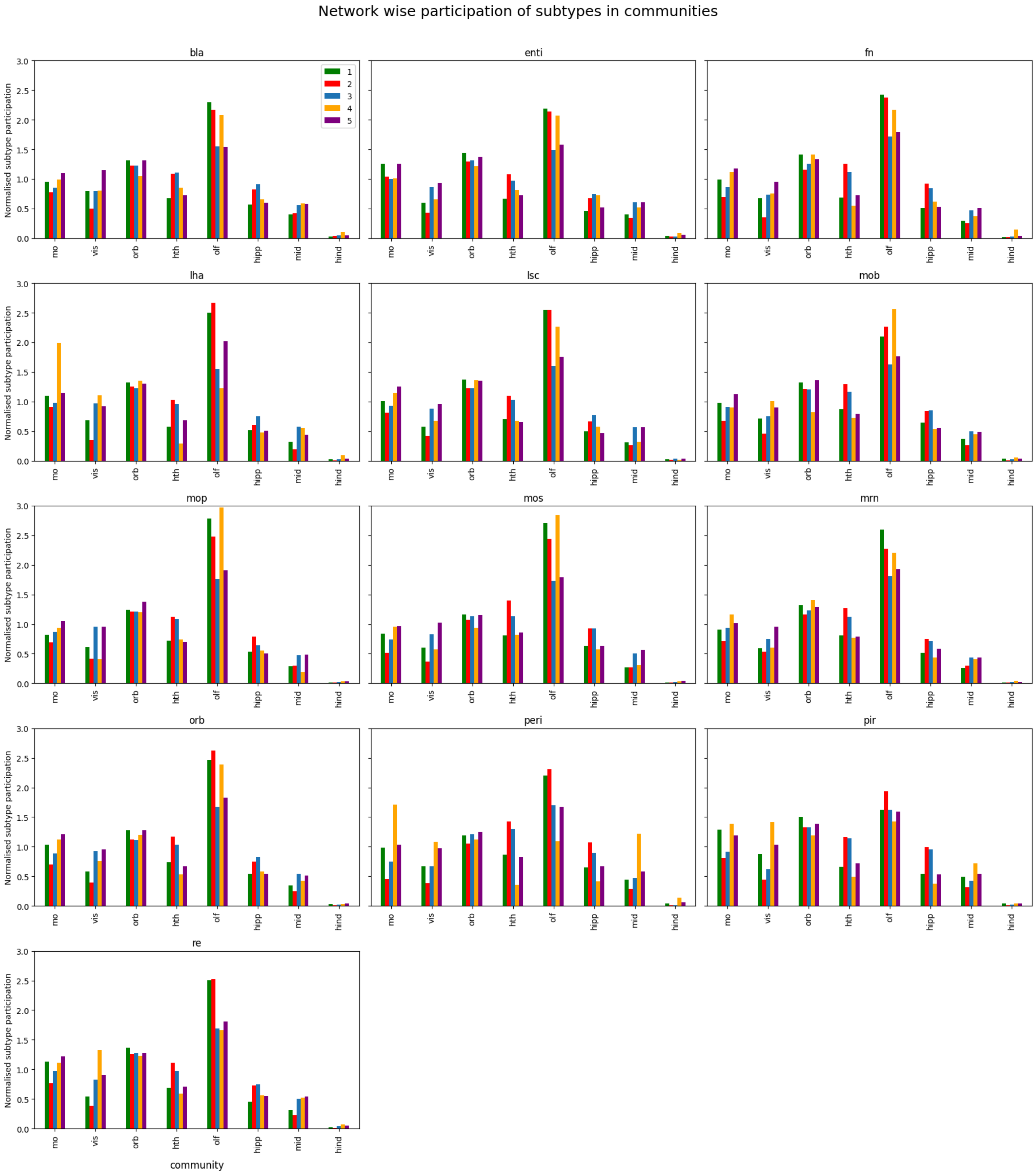


Community participation in individual subtypes across all networks

Fig. S5.1: Biexponential fit for partial FPT

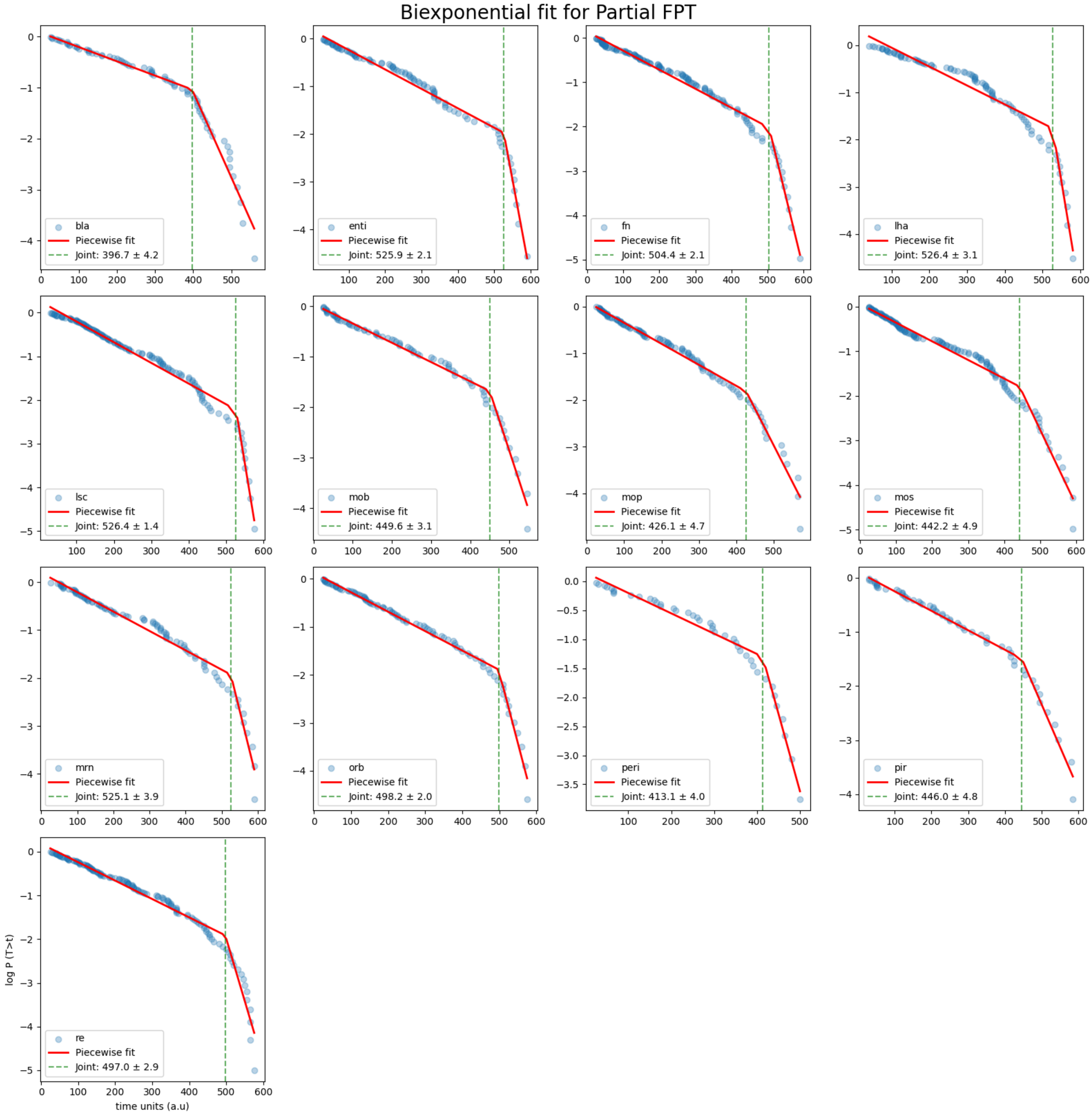

Fig. S5.2: Biexponential fit for full FPT

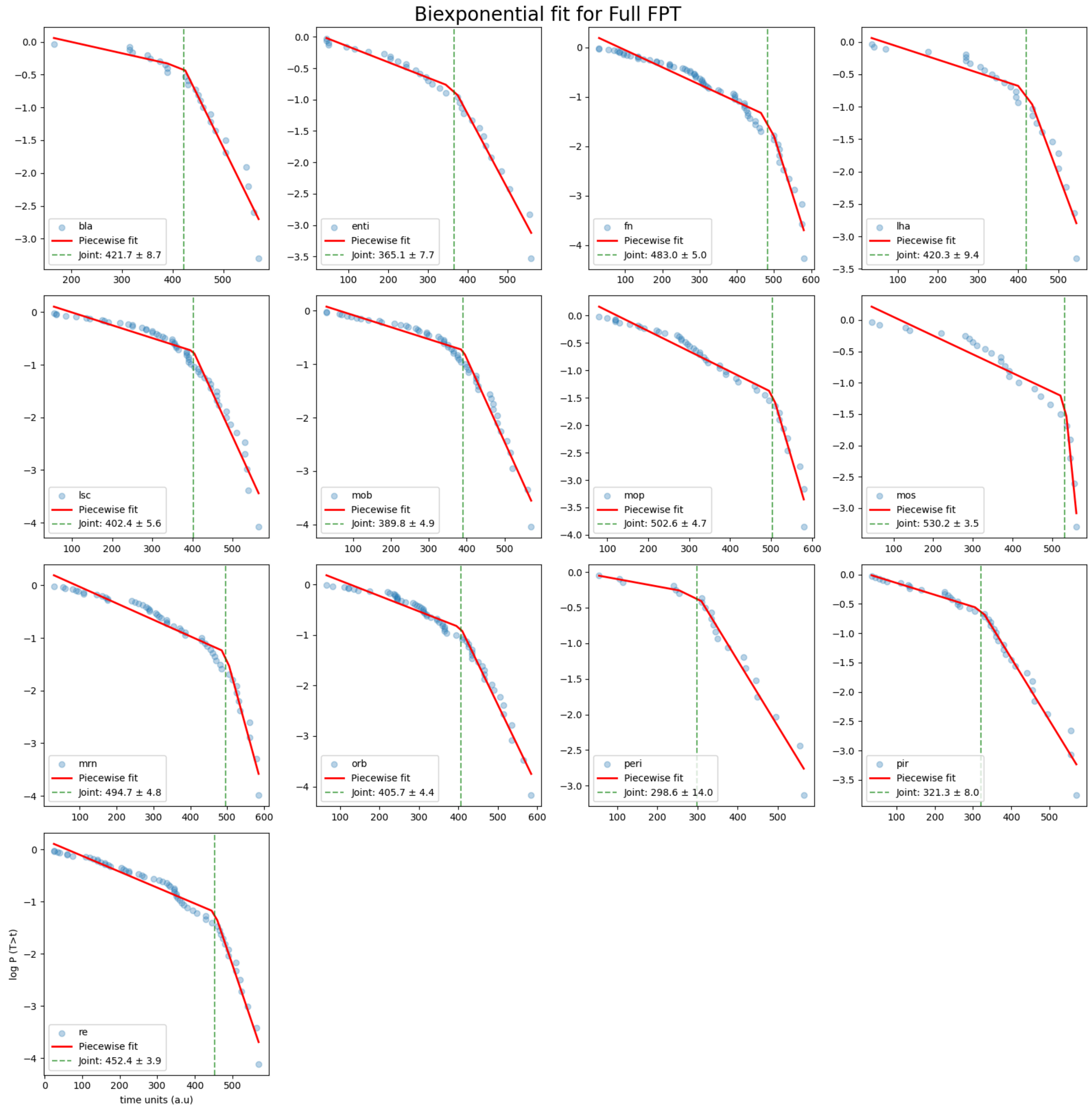

Fig. S5.3: Biexponential fit for part ST

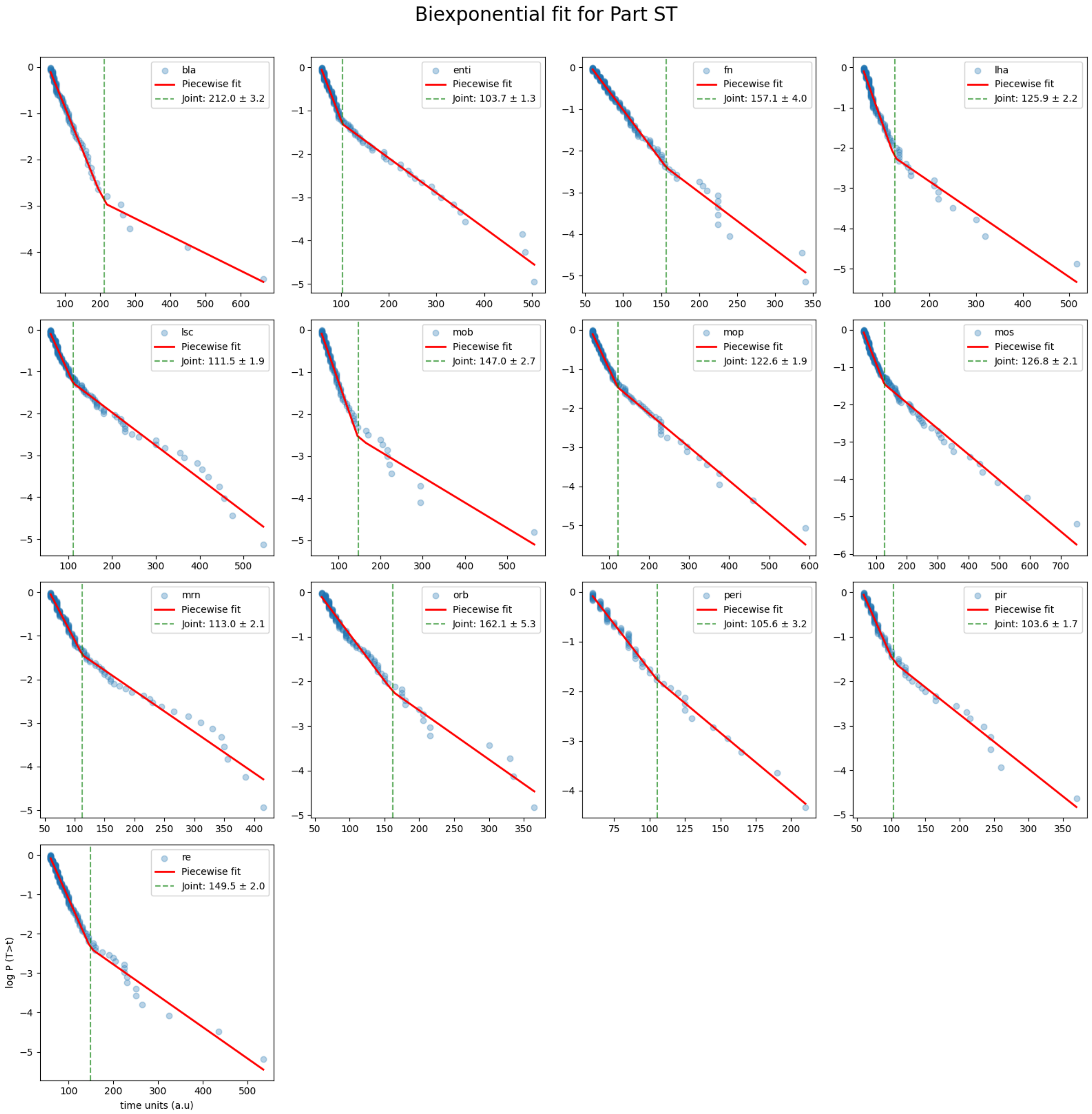

Fig. S5.4: Biexponential fit for full ST

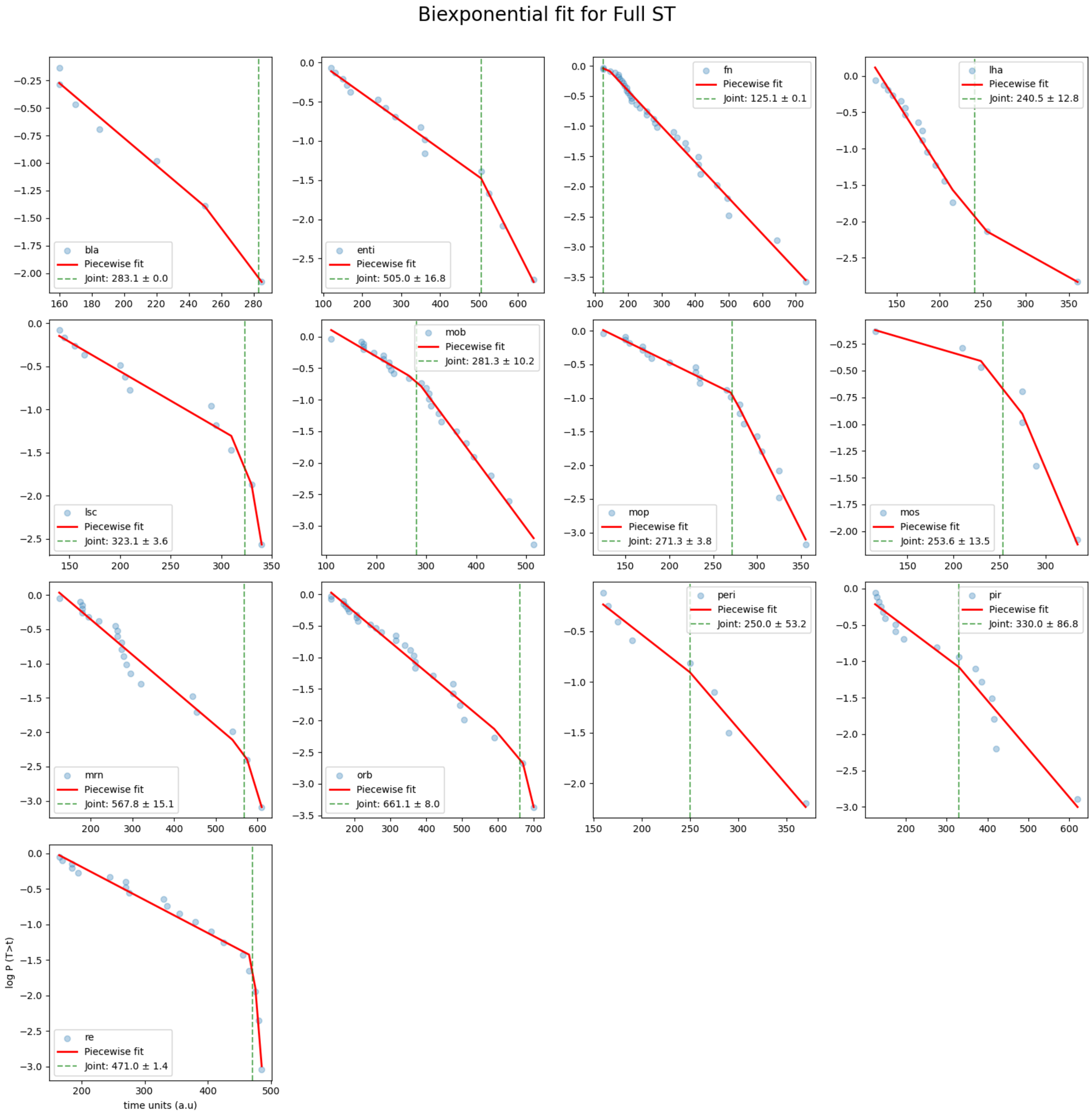

Fig. S5.5 Clusterwise full transitions ST biexponential fit

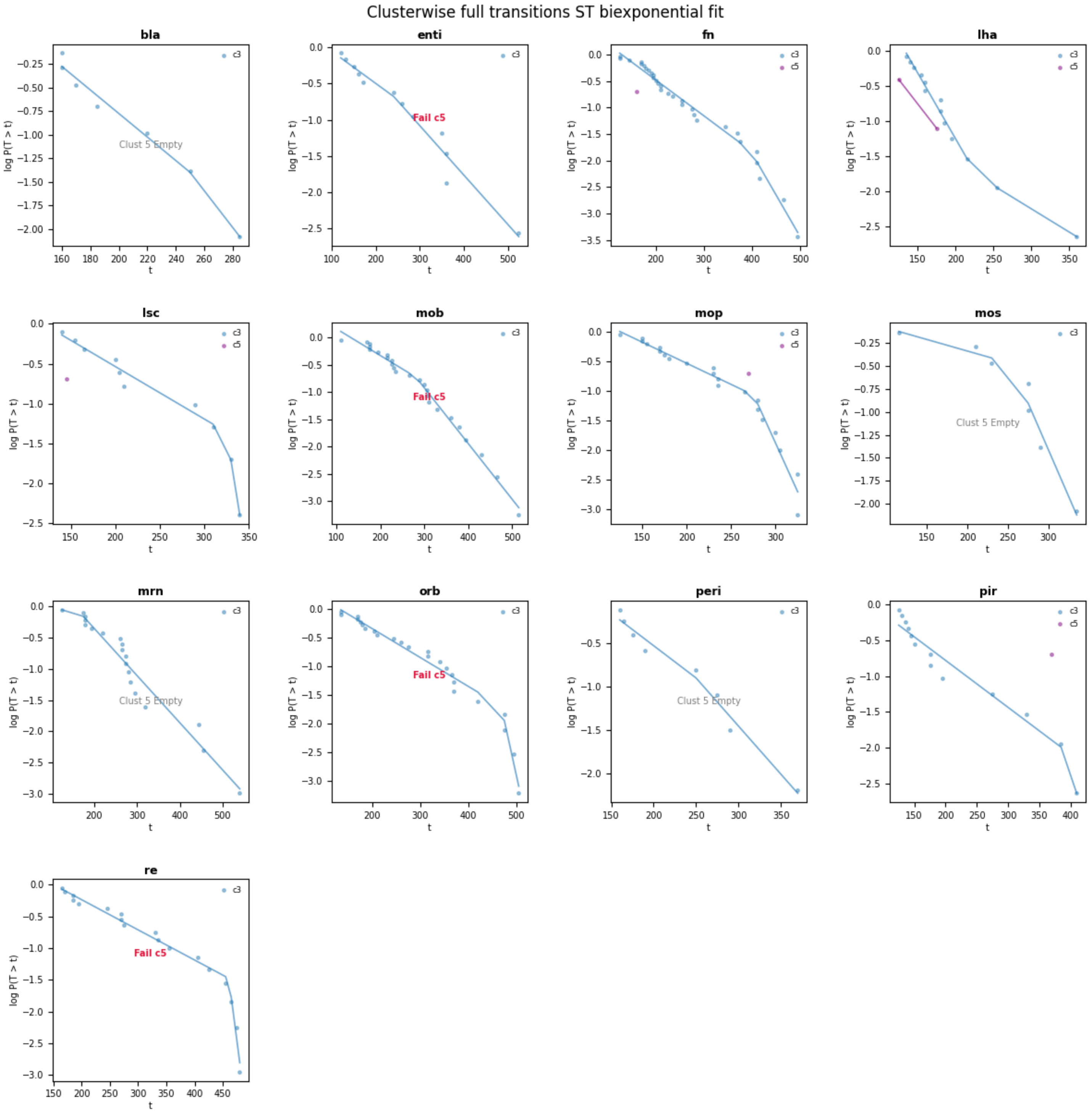

### Fig. S5.6 - Clusterwise full transitions FPT biexponential fit

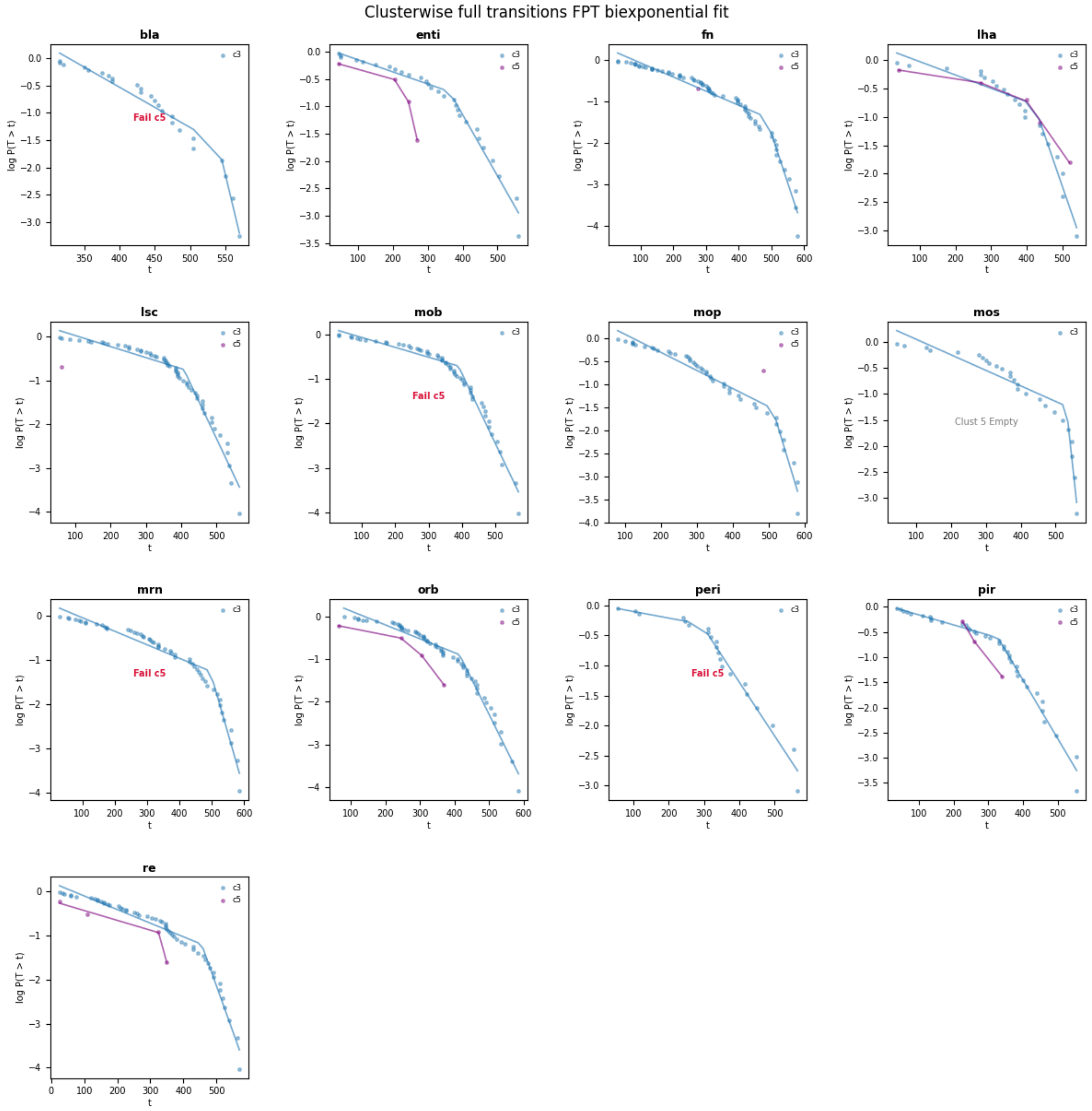

Fig. S5.7-Clusterwise full transitions FPT biexponential fit

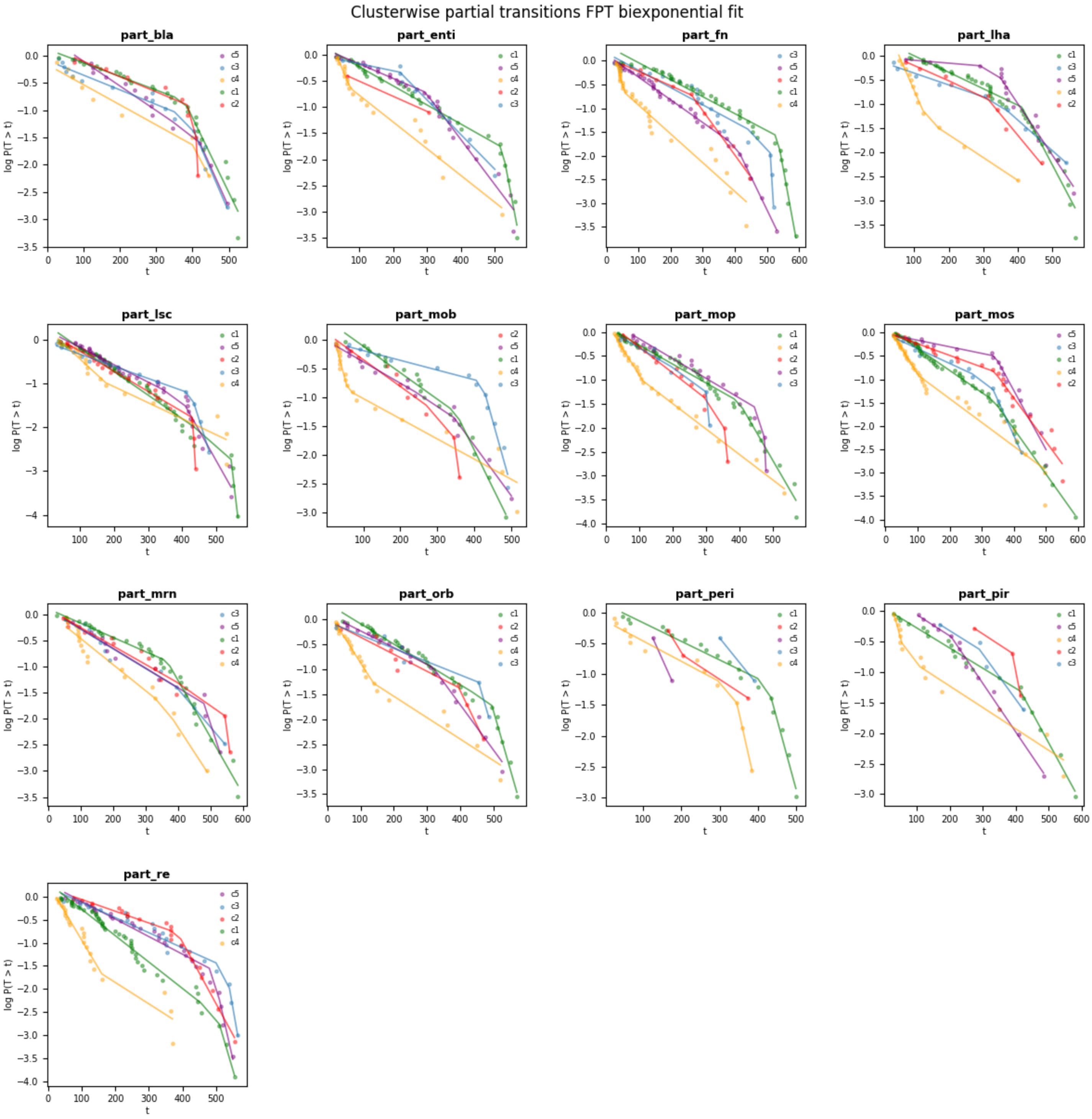


**Table 5.1: Pairwise comparison of each subtype for a specific parameter**

**Pairwise Significance Matrix for Transition Time**

| Comparison | Raw p-value | Corrected p-value | Significance |
|---|---|---|---|
| Subtype 1 vs Subtype 2 | 4.42e-01 | 1.00e+00 | ns |
| Subtype 1 vs Subtype 3 | 1.65e-05 | < 0.001 | <0.001 |
| Subtype 1 vs Subtype 4 | 6.08e-01 | 1.00e+00 | ns |
| Subtype 1 vs Subtype 5 | 4.02e-04 | 2.41e-03 | <0.01 |
| Subtype 2 vs Subtype 3 | 1.65e-05 | < 0.001 | <0.001 |
| Subtype 2 vs Subtype 4 | 9.59e-01 | 1.00e+00 | ns |
| Subtype 2 vs Subtype 5 | 3.47e-03 | 1.73e-02 | <0.05 |
| Subtype 3 vs Subtype 4 | 1.65e-05 | < 0.001 | <0.001 |
| Subtype 3 vs Subtype 5 | 1.65e-05 | < 0.001 | <0.001 |
| Subtype 4 vs Subtype 5 | 2.10e-02 | 8.41e-02 | ns |

**Pairwise Significance Matrix for FPT**

| Comparison | Raw p-value | Corrected p-value | Significance |
|---|---|---|---|
| Subtype 1 vs Subtype 2 | 3.05e-01 | 1.00e+00 | ns |
| Subtype 1 vs Subtype 3 | 1.00e+00 | 1.00e+00 | ns |
| Subtype 1 vs Subtype 4 | 1.20e-04 | < 0.001 | <0.001 |
| Subtype 1 vs Subtype 5 | 4.02e-02 | 2.41e-01 | ns |
| Subtype 2 vs Subtype 3 | 3.83e-01 | 1.00e+00 | ns |
| Subtype 2 vs Subtype 4 | 5.09e-05 | < 0.001 | <0.001 |
| Subtype 2 vs Subtype 5 | 3.30e-01 | 1.00e+00 | ns |
| Subtype 3 vs Subtype 4 | 1.48e-04 | 1.03e-03 | <0.01 |
| Subtype 3 vs Subtype 5 | 6.49e-02 | 3.24e-01 | ns |
| Subtype 4 vs Subtype 5 | 2.61e-05 | < 0.001 | <0.001 |

**Pairwise Significance Matrix for COV**

| Comparison | Raw p-value | Corrected p-value | Significance |
|---|---|---|---|
| Subtype 1 vs Subtype 2 | 1.65e-05 | < 0.001 | <0.001 |
| Subtype 1 vs Subtype 3 | 1.65e-05 | < 0.001 | <0.001 |
| Subtype 1 vs Subtype 4 | 1.65e-05 | < 0.001 | <0.001 |
| Subtype 1 vs Subtype 5 | 1.65e-05 | < 0.001 | <0.001 |
| Subtype 2 vs Subtype 3 | 1.65e-05 | < 0.001 | <0.001 |
| Subtype 2 vs Subtype 4 | 1.65e-05 | < 0.001 | <0.001 |
| Subtype 2 vs Subtype 5 | 8.58e-04 | < 0.001 | <0.001 |
| Subtype 3 vs Subtype 4 | 1.65e-05 | < 0.001 | <0.001 |
| Subtype 3 vs Subtype 5 | 1.65e-05 | < 0.001 | <0.001 |
| Subtype 4 vs Subtype 5 | 1.65e-05 | < 0.001 | <0.001 |

**Pairwise Significance Matrix for ST**

| Comparison | Raw p-value | Corrected p-value | Significance |
|---|---|---|---|
| Subtype 1 vs Subtype 2 | 2.93e-01 | 1.00e+00 | ns |
| Subtype 1 vs Subtype 3 | 2.61e-05 | < 0.001 | <0.001 |
| Subtype 1 vs Subtype 4 | 7.20e-01 | 1.00e+00 | ns |
| Subtype 1 vs Subtype 5 | 5.38e-01 | 1.00e+00 | ns |
| Subtype 2 vs Subtype 3 | 4.08e-03 | 2.86e-02 | <0.05 |
| Subtype 2 vs Subtype 4 | 4.12e-01 | 1.00e+00 | ns |
| Subtype 2 vs Subtype 5 | 4.42e-01 | 1.00e+00 | ns |
| Subtype 3 vs Subtype 4 | 4.09e-05 | < 0.001 | <0.001 |
| Subtype 3 vs Subtype 5 | 5.09e-05 | < 0.001 | <0.001 |
| Subtype 4 vs Subtype 5 | 7.00e-01 | 1.00e+00 | ns |

**Table for 5F: Pairwise comparison of each subtype for Time Inertia**

**Pairwise Significance: Partial FPT Time Inertia**

| Comparison | Raw p-value | Corrected p-value | Significance |
|---|---|---|---|
| Cluster c0 vs Cluster c1 | 2.74e-02 | 1.64e-01 | ns |
| Cluster c0 vs Cluster c2 | 1.12e-01 | 4.48e-01 | ns |
| Cluster c0 vs Cluster c3 | 1.33e-04 | 1.33e-03 | <0.01 |
| Cluster c0 vs Cluster c4 | 2.74e-02 | 1.64e-01 | ns |
| Cluster c1 vs Cluster c2 | 8.58e-01 | 1.00e+00 | ns |
| Cluster c1 vs Cluster c3 | 1.48e-03 | 1.33e-02 | <0.05 |
| Cluster c1 vs Cluster c4 | 9.59e-01 | 1.00e+00 | ns |
| Cluster c2 vs Cluster c3 | 5.62e-03 | 3.93e-02 | <0.05 |
| Cluster c2 vs Cluster c4 | 8.37e-01 | 1.00e+00 | ns |
| Cluster c3 vs Cluster c4 | 2.09e-03 | 1.67e-02 | <0.05 |

**Pairwise Significance: Full / Partial FPT Time Inertia**

| Comparison | U-statistic | p-value | Significance |
|---|---|---|---|
| Subtype 3 (Partial) vs Subtype 3 (Full) | 54.50 | 1.30e-01 | ns |
| Subtype 3 (Partial) vs Subtype 5 (Full) | 74.00 | 1.21e-01 | ns |
| Subtype 5 (Partial) vs Subtype 3 (Full) | 46.00 | 5.12e-02 | ns |
| Subtype 5 (Partial) vs Subtype 5 (Full) | 71.00 | 1.85e-01 | ns |

**Table for 5H: Pairwise comparison of each subtype for Time Inertia**

| Comparison | U-Statistic | Raw p-value | Corrected | p-value Significance |
|---|---|---|---|---|
| Full vs Partial (EARLY) | 155.0 | 3.17e-05 | < 0.001 | <0.001 |
| Full vs Partial (LATE) | 48.0 | 1.09e-01 | 1.09e-01 | ns |
| Early vs Late (FULL) | 131.0 | 7.31e-04 | 1.46e-03 | p<0.01 |
| Early vs Late (PARTIAL) | 5.0 | 5.09e-05 | < 0.001 | p<0.001 |

**Table for 5 I, J and 6 (T5.1):**

| nw / p-val | bla | enti | lha | lsc | mob | mop | mos | mrn | orb | peri | pir | re |
|---|---|---|---|---|---|---|---|---|---|---|---|---|
| FPT λ early (P) | 0.0000e+00 | 1.8408e-01 | 2.2647e-01 | 1.3490e-05 | 4.9758e-05 | 2.0496e-01 | 9.5306e-01 | 2.0496e-01 | 3.2617e-02 | 3.1207e-05 | 1.5442e-08 | 9.5306e-01 |
| FPT λ late (P) | 0.0000e+00 | 3.6377e-06 | 1.1379e-09 | 0.0000e+00 | 0.0000e+00 | 0.0000e+00 | 0.0000e+00 | 7.9511e-02 | 3.2342e-04 | 1.5320e-06 | 0.0000e+00 | 1.0517e-04 |
| FPT λ early (F) | 7.4760e-02 | 7.8493e-04 | 8.4555e-04 | 1.2199e-03 | 2.8130e-06 | 7.4808e-01 | 4.6282e-01 | 4.6282e-01 | 3.3738e-01 | 7.9422e-07 | 1.3159e-06 | 1.8017e-01 |
| FPT λ late (F) | 2.8421e-10 | 0.0000e+00 | 1.1546e-06 | 3.4100e-10 | 6.4393e-14 | 7.2673e-01 | 3.2543e-12 | 6.2258e-01 | 1.4648e-12 | 0.0000e+00 | 0.0000e+00 | 1.0286e-01 |
| ST λ early (P) | 0.0000e+00 | 0.0000e+00 | 0.0000e+00 | 5.8487e-03 | 0.0000e+00 | 3.2914e-10 | 0.0000e+00 | 1.1237e-03 | 1.3323e-15 | 0.0000e+00 | 0.0000e+00 | 8.0274e-04 |
| ST λ late (P) | 0.0000e+00 | 0.0000e+00 | 0.0000e+00 | 0.0000e+00 | 0.0000e+00 | 0.0000e+00 | 0.0000e+00 | 0.0000e+00 | 5.7588e-08 | 0.0000e+00 | 3.9088e-04 | 0.0000e+00 |
| ST T stabilisation time (P) | 0.0000e+00 | 0.0000e+00 | 4.4674e-11 | 0.0000e+00 | 1.4409e-01 | 4.5075e-14 | 9.8921e-11 | 0.0000e+00 | 8.7838e-01 | 0.0000e+00 | 0.0000e+00 | 2.6876e-01 |
| NO (F) | 2.4345e-04 | 2.1149e-03 | 1.0963e-03 | 1.0000e+00 | 1.0000e+00 | 1.1443e-01 | 3.3069e-06 | 3.7560e-01 | 1.0000e+00 | 4.2219e-05 | 3.9641e-02 | 1.0000e+00 |
| NO (P) | 2.9919e-05 | 2.5071e-01 | 6.5206e-02 | 1.0000e+00 | 1.8375e-02 | 8.8092e-01 | 1.0000e+00 | 1.9946e-01 | 3.6022e-02 | 2.2602e-10 | 7.1609e-05 | 1.0000e+00 |
| TT (P) | 1.0000e+00 | 1.0000e+00 | 4.0675e-01 | 1.000000 | 1.000000 | 4.0675e-01 | 4.0675e-01 | 1.000000 | 1.000000 | 1.3087e-01 | 1.000000 | 1.000000 |
| TT (F) | 1.000000 | 1.000000 | 0.443728 | 1.000000 | 1.000000 | 0.443728 | 0.443728 | 1.000000 | 1.000000 | 0.141772 | 1.000000 | 1.000000 |
| COV (P) | 1.0000e+00 | 1.0000e+00 | 1.0000e+00 | 1.0000e+00 | 1.0000e+00 | 4.5612e-01 | 1.0000e+00 | 1.0000e+00 | 2.8943e-01 | 1.0000e+00 | 1.0000e+00 | 1.0000e+00 |
| COV (F) | 0.289324 | 1.000000 | 0.007615 | 1.000000 | 1.000000 | 0.617734 | 1.000000 | 0.007235 | 0.106112 | 1.000000 | 1.000000 | 1.000000 |
